# The molecular basis of lamin-specific chromatin interactions

**DOI:** 10.1101/2024.08.05.604734

**Authors:** Baihui Wang, Rafael Kronenberg-Tenga, Valentina Rosti, Emanuele Di Patrizio Soldateschi, Qiang Luo, Louise Pinet, Matthias Eibauer, Rajaa Boujemaa-Paterski, Benjamin Schuler, Chiara Lanzuolo, Ohad Medalia

**Affiliations:** Department of Biochemistry, University of Zurich, Winterthurerstrasse 190, 8057 Zurich, Switzerland; Istituto Nazionale Genetica Molecolare “Romeo ed Enrica Invernizzi”, Milan, Italy; Institute of Biomedical Technologies, National Research Council, Milan, Italy

## Abstract

In the cell nucleus, chromatin is anchored to the nuclear lamina, a network of lamin filaments and binding proteins that underly the inner nuclear membrane. The nuclear lamina is involved in chromatin organisation through the interaction of lamina-associated domains (LADs) within the densely packed heterochromatin regions. Employing cryo-focused ion beam (cryo-FIB) milling in conjunction with cryo-electron tomography (cryo-ET), we analysed the distribution of nucleosomes at the lamin-chromatin interface. Depletion of lamin A/C reduced the concentration of nucleosomes at the nuclear periphery, suggesting that lamins are directly involved in the interaction with chromatin. Using cryo-electron microscopy (cryo-EM), we then identified the specific binding motif of the lamin A tail domain that interacts with nucleosomes, distinguishing it from the other lamin isoforms. Furthermore, we examined chromatin structure dynamics using a genome-wide analysis that revealed lamin-dependent macroscopic-scale alterations in gene expression and chromatin remodelling. Our findings provide detailed insights into the dynamic and structural interplay between lamin isoforms and chromatin, molecular interactions which are shaping chromatin architecture and epigenetic regulation.

## Main Text

Nuclear lamins provide mechanical support and safeguard the integrity of the cell nucleus. At the nuclear lamina (NL), lamins contribute to the organisation of chromatin^1, 2^. In mammals, four main lamin isoforms are expressed: the A-type lamins (A and C) and the B-type lamins (B1 and B2)^3^. During their posttranslational processing, lamin A, B1, and B2 are farnesylated at their C-terminus. Notably, while lamin A loses its farnesyl group through a subsequent cleavage, the B-type lamins retain their farnesylation, serving as anchors to the inner nuclear membrane (INM)^4^. The two types of lamin proteins have been described to form two separate but interconnected filamentous meshworks^5^, wherein the B-type lamin meshwork is localised closer to the nuclear membrane^6,7^. Previous investigations of the nuclear lamina of mouse embryonic fibroblasts (MEFs) have shown a 14-16 nm thick meshwork consisting of ∼3.5 nm thick lamin filaments^8, 9^.

Knock-out of A-type lamins in MEFs significantly impacts nuclear mechanics by reducing stiffness, structural integrity^10–12^, and decoupling nuclear and cytoplasmic forces^13^. These observations suggest direct interactions between lamin A/C and chromatin^14^ at the docking sites of heterochromatic regions^15–18^. These large sites, commonly termed LADs, can extend up to 10 Mb in length and are typically enriched with the H3K9me3 repressive mark, rendering them transcriptionally inactive. However, 10% of the genes inside LADs are still expressed^19^. Interestingly, A-type lamins were shown to also interact with euchromatic DNA^20^ and with facultative H3K27me3 enriched chromatin^21–24^, while lamin B1 is also localised to open and actively expressed chromatin regions^25^. Insufficient lamin expression results in genome remodelling, manifested by alterations in histone markers and the detachment of LADs^26, 27^. Genome-wide approaches such as HiC or DamID have detected lamin-dependent chromatin alterations^17, 26, 27^; however, the direct molecular interactions and effects of the lamin isoforms on chromatin organisation remained elusive.

## Results

### Chromatin and lamin organisation at the nuclear envelope imaged by cryo-electron tomography

To dissect the contribution of nuclear lamins to the organisation of the nuclear envelope (NE), we analysed tomograms of cryo-FIB milled, vitrified cells (Methods section), in which the components of the NE were observable, e.g., nuclear lamins, nuclear pore complexes (NPCs), and nucleosomes (Fig. 1a, Extended Data Fig. 1a), and their coordinates were extracted from each nuclear volume. Subsequently, the nucleosomes were subjected to subtomogram averaging^28^, yielding a 14 Å resolved structure exhibiting the typical appearance in which a DNA double-strand is wrapped around a histone octamer (Extended Data Fig. 2a). We then measured the in situ local concentrations and distributions of lamins and nucleosomes (Fig. 1). LADs, where chromatin is attached to the nuclear lamina, are commonly attributed as heterochromatin-rich domains^29^. Our analysis resolved varying nucleosome concentrations within these regions (Fig. 1a), showing a fine modulation within both high concentration values (Fig. 1a, upper right) as well as lower nucleosome concentrations (Fig. 1a, lower right, Extended Data Fig. 2b), in line with limited presence of open chromatin regions near NPCs^30^ or within LADs^31^.

**Fig. 1.**
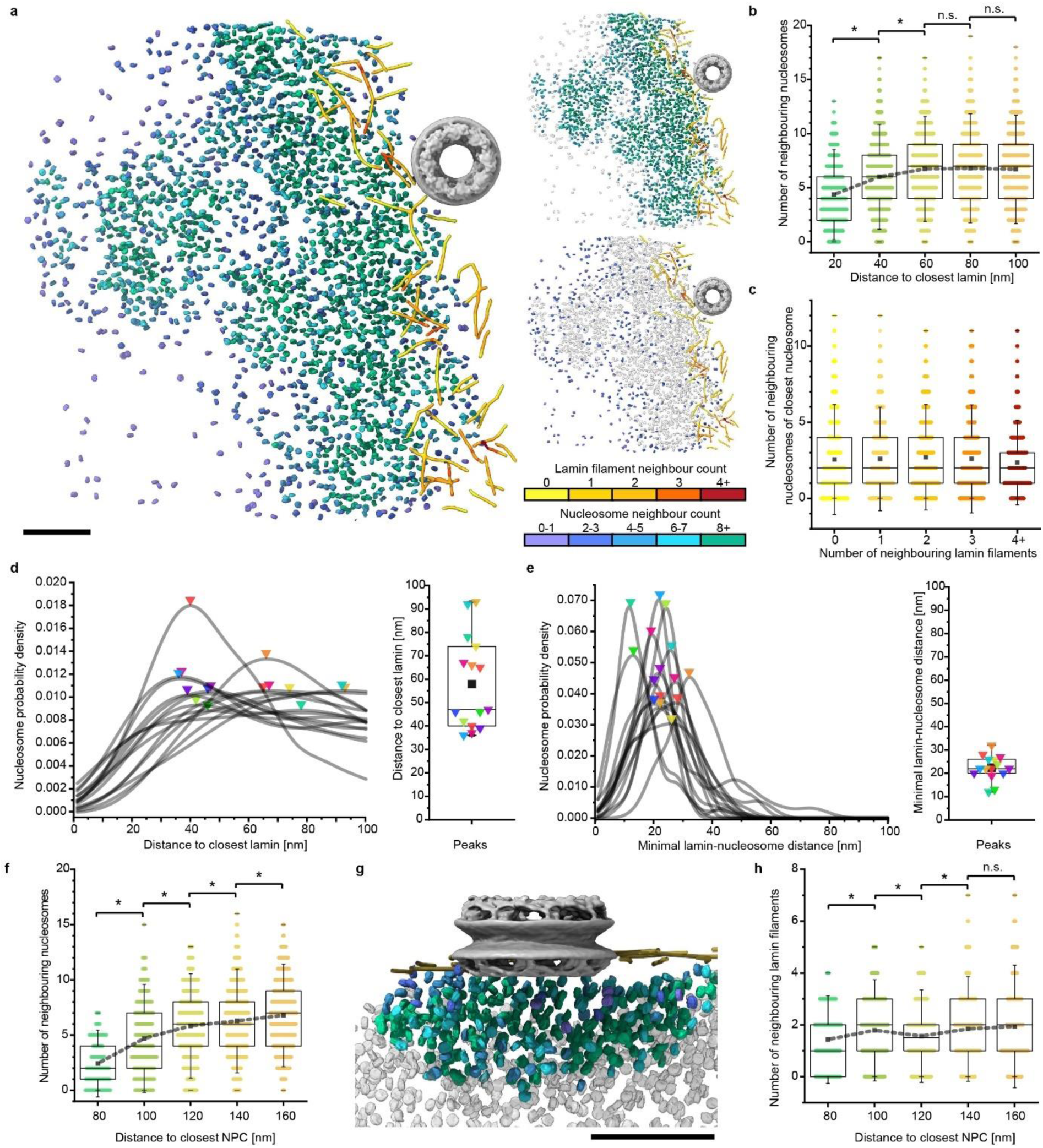
The local concentration and distribution of nucleosomes at the nuclear envelope. **a.** Segmented view of a tomogram depicting the nuclear envelope of a MEF cell. Nucleosomes and lamins are coloured based on their local concentration, from purple to green and yellow to red, respectively. At the position of an NPC, its structure was manually placed (grey, EMDB: 12814). The image was split into two, depicting local nucleosome concentrations, either with 4 or more neighbours (upper right panel) or nucleosomes with 3 or fewer neighbours in grey (lower right panel). **b.** The local concentration of nucleosomes is shown as the number of neighbouring nucleosomes within a distance of 24 nm as a function of their distance from the NL. The nucleosome concentration increased with increasing distance from the NL. Further than 60 nm away from the NL, the nucleosome concentration reached a plateau of 7 neighbours in average, which corresponds to a concentration of 1.4x10^-4^ nucleosomes/nm^3^. The average lamin-independent nucleosome concentration was measured in **Extended Data** Fig. 2d. **c.** For each lamin segment, the closest nucleosome was identified, and the count of nucleosome neighbours was plotted against the number of lamin filaments from which the segments originated. The nucleosome concentration was consistently low, with a median of 2 neighbours close to the NL, independent of the filament concentration. This indicates that there is no apparent linear correlation between lamin and nucleosome concentration. **d.** For each tomogram, the nucleosome concentration probability was plotted as a function of the distance to the closest lamin filament in each tomogram, using a line fit for an average of ∼2400 particles per tomogram. High-density peaks are indicated by triangles. Their associated distance of peaks was then plotted (right panel), indicating a median value of 47 nm. **e.** The nucleosome probability density as a function of the distance between nucleosomes and lamins was plotted for each tomogram, i.e., the minimal distance between lamin segments and their closest nucleosomes. The peaks are indicated by triangles and were plotted against associated distance values (right panel). The median distance was 22 ± 5 nm. **f.** Similarly to **b**, the nucleosome concentration was plotted against the distance to the centre of the closest NPC. The further away from an NPC, the steadier the increase in nucleosome concentration. **g.** Segmented view of a measurement in **f**. The nucleosomes are coloured based on their concentration, as in **a,** up to 160 nm away from the centre of the NPC. The nucleosomes are presented in grey for longer distances. Lamins are shown in dark yellow. **h**. The concentration of lamin filaments was plotted against their distance to the closest NPC. Scale bar 100 nm. All boxplots show a box between the 25^th^ and 75^th^ percentiles, the median as a horizontal line, the mean as a black square, and whiskers represent 1.5 standard deviations. Significance was calculated using a one-way ANOVA with a significance level of 0.05. * = p < 0.05, n.s. = not significant.

To study the interplay between lamins and chromatin, we measured the local concentration of nucleosomes as a function of the distance from their closest lamin filament (Fig. 1b), focusing our analysis on a 100 nm wide region closest to the NE. In wild-type (WT) MEFs, this measurement revealed a specific nucleosome pattern, which shows a gradual increase in nucleosome concentration between ∼20 to ∼60 nm distance from the lamin filaments, followed by a constant nucleosome concentration further away. This suggests a less dense chromatin organisation at the NL-chromatin interface, where LADs interact directly with the lamina, than 60-100 nm away from the NL. Interestingly, the concentration of nucleosomes is not linearly correlated with the local concentration of lamin filaments at the NL (Fig. 1c).

Next, we measured the position of individual nucleosomes as a function of their distance from the NL (i.e., the closest lamin filament) in every reconstructed nuclear volume (Fig. 1d, left). In half of the nuclear volumes, the highest nucleosome concentration was located within a narrow range of 35-47 nm from the NL (Fig 1d, right) and remained at similar levels for the next ∼50 nm. We also measured the distance between lamins and their closest nucleosome (Fig. 1e, left), finding an average distance of 22±5 nm across all tomograms (Fig. 1e, right). This distance would allow a possible interaction between nucleosomes and the lamins via lamins’ elongated tail domains^14^.

A previous study has described that chromatin is excluded from the vicinity of NPCs^32^. We investigated the concentration of nucleosomes around NPCs and its correlation with the lamin meshwork density. To this end, we determined the coordinates of NPCs and measured the concentration of nucleosomes as a function of the distance to the centre of the closest NPC (Fig. 1f, g). Given that the diameter of NPCs is ∼110 nm in MEFs^33^, we measured the local concentration of nucleosomes within a radial distance of 60 to 160 nm from the NPCs’ central coordinates. The nucleosome concentration increases with increasing distance from the NPCs, reaching the previously measured maximal values (Fig. 1b) only at a distance of >150 nm. However, the concentration of lamin filaments is less affected by distance from NPCs (Fig. 1h). Conclusively, lamin filaments are affected by specific NPC components^34^; however, their concentration is steady around NPCs.

### The effect of lamin isoforms on the nuclear envelope

To resolve the effect of A- and B-type lamins on chromatin organisation at the nanometre scale, we subjected *Lmna*^-/-^ (LmnaKO) and *Lmnb1*^-/-^+*Lmnb2*^-/-^ (LBDKO) MEFs to cryo-FIB milling, cryo-ET, and image analysis, as described for WT MEFs. Lamins were segmented from 13 LmnaKO and 15 LBDKO MEF tomograms, and the local concentrations of lamin filaments were determined (Fig. 2a, Extended Data Fig. 1b, c). While the lamin meshworks underlying the NE appeared visually similar, substantial differences were observed in the proportion of single lamin filaments without neighbouring filaments. In WT MEFs, the majority of lamin filaments were organised into a closely packed meshwork, with only 16% of isolated filaments (Fig. 2a). In the absence of A-type lamins, up to 28% of filaments were isolated, whereas the removal of B-type lamins resulted in a modest increase to 19% of lamins lacking closely neighbouring filaments (Fig. 2a). An additional effect of lamin A/C knockout was the overall decrease in the mean count of neighbouring filaments, from 2.1 in WT to 1.6 in LmnaKO cells (Extended Data Fig. 2c). Interestingly, the lamin meshwork density in LBDKO cells was not substantially altered (Extended Data Fig. 2c), possibly reflecting the enrichment of A-type lamins over B-types in these cells.

**Figure 2.**
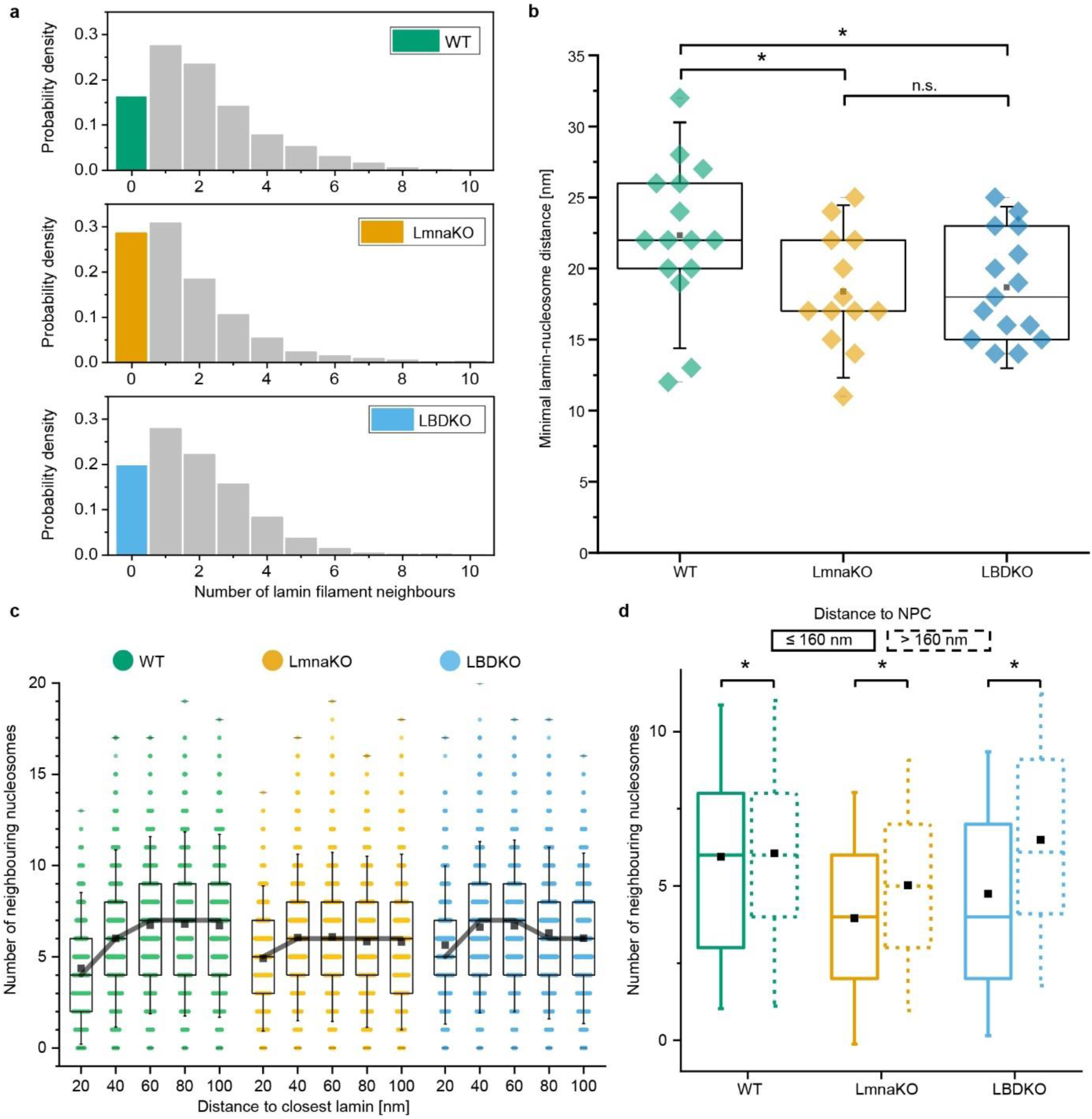
The alteration of filament and nucleosome distribution in lamin knockout cells. **a.** The lamin meshwork density of filaments is shown. The number of neighbouring filaments detected at a distance of <24 nm around each lamin segment was plotted. The histograms indicated the different cell lines. The fraction of lamins without neighbouring filament was significantly increased from 16% in the WT to 28% in LmnaKO cells, to 19% in LBDKO cells, with a drop of the mean neighbour count of 2.1, 1.6 and 1.9 in each cell line, respectively (see **Extended Data** Fig. 2c). **b.** The overall minimal lamin-to-nucleosome distances were plotted for each cell line (as in **Fig. 1e**). Each datapoint represents the highest nucleosome density distance per tomogram. The median of the minimal lamin-to-nucleosome distances were 18.4 ± 4.1 nm for LmnaKO cells and 18.7 ± 3.8 nm for LBDKO cells, corresponding to ∼82% of the WT’s mean. **c.** The concentration of nucleosomes as a function of their distance from the NL, plotted as in **Fig. 1b**. The median concentration of nucleosomes for each distance was connected by a black line. Changes in the nucleosome concentration profile between knockout cells and WT were detected. Lamin-independent average nucleosome concentration was measured in **Extended Data** Fig. 2d. **d.** For each cell line, nucleosomes were grouped into two groups by their distance to the centre of the closest NPC of ≤ 160 nm (solid lines) or >160 nm (dashed lines). In the WT, there was no significant correlation between the concentration of nucleosomes in the two groups. However, in both knockout cells, the nucleosome concentration is lower at the NPC proximal 160 nm. All boxplots show a box between the 25^th^ and 75^th^ percentiles, the median as a horizontal line, the mean as a black square, and whiskers represent 1.5 standard deviations. Significance was calculated using a one-way ANOVA with a significance level of 0.05. * = p < 0.05, n.s. = not significant.

The decrease in the minimal distance between lamins and nucleosomes demonstrated that nucleosomes were positioned closer to the NL in nuclei lacking one of the lamin types (Fig. 2b). This may be attributed to the lower occupancy of lamins and their associated factors at the NE in lamin knock-out cell lines. Surprisingly, the overall nucleosome concentration was significantly reduced in LmnaKO cells at the nuclear periphery while remaining unchanged in LBDKO cells (Fig. 2c, Extended Data Fig. 2d). In LmnaKO cells, nucleosome concentrations peaked at a shorter distance of ∼40 nm from the NL and did not reach the same concentration level as observed in WT cells (Fig. 2c). Noteworthily, LBDKO cells exhibited a distinct gradient of nucleosome concentration throughout the 20-60 nm layer away from the NL, with a slight reduction observed further away. These findings indicate that predominantly A-type lamins contribute to the precise chromatin organisation at the NE and may be involved in direct molecular lamin-chromatin interactions.

Moreover, in both knock-out cell lines, we noted a statistically significant alteration in the concentration of nucleosomes around NPCs. Nucleosomes were less concentrated in the immediate vicinity of the NPC (≤ 160 nm away from the centre coordinates of NPCs, Fig. 2d) compared to WT, indicating that alterations in lamins affect the chromatin organisation around NPCs as well.

### Lamin A interacts with H2A-H2B and nucleosomes via its tail domain

Previous studies have shown the possibility of a direct molecular interaction between chromatin and lamins^35, 36^ via their tail domains^37–39^. Based on sequence alignment (Fig. 3a), we designed a set of lamin A tail truncations (Extended Data Fig. 3a) and studied their binding properties to H2A-H2B heterodimers and to reconstituted nucleosomes (Fig. 3b, c). Using pull-down assays (Methods section), we found that the C-terminal tail of lamin A (LA 394-646) effectively binds to H2A-H2B heterodimers (Extended Data Fig. 3b). Notably, a truncated tail domain spanning from amino acid 430 to 585 (LA 430-585) exhibited a similar binding property as the full-length tail domain (LA 394-646), indicating that this region is sufficient for binding the H2A-H2B heterodimers. In contrast, truncated tail regions comprising either the NLS and the Ig-like domain (LA 394-548) or only the Ig-like domain (LA 430-560) did not bind to H2A-H2B heterodimers, although weak binding was detected with LA 430-579 (Extended Data Fig. 3b). Fluorescence correlation spectroscopy (FCS) measurements indicated a dissociation constant K_D_ of 12 ± 1 µM for the binding of LA 430-585 to purified H2A-H2B heterodimers (Fig. 3b, Extended Data Fig. 3c).

**Figure 3.**
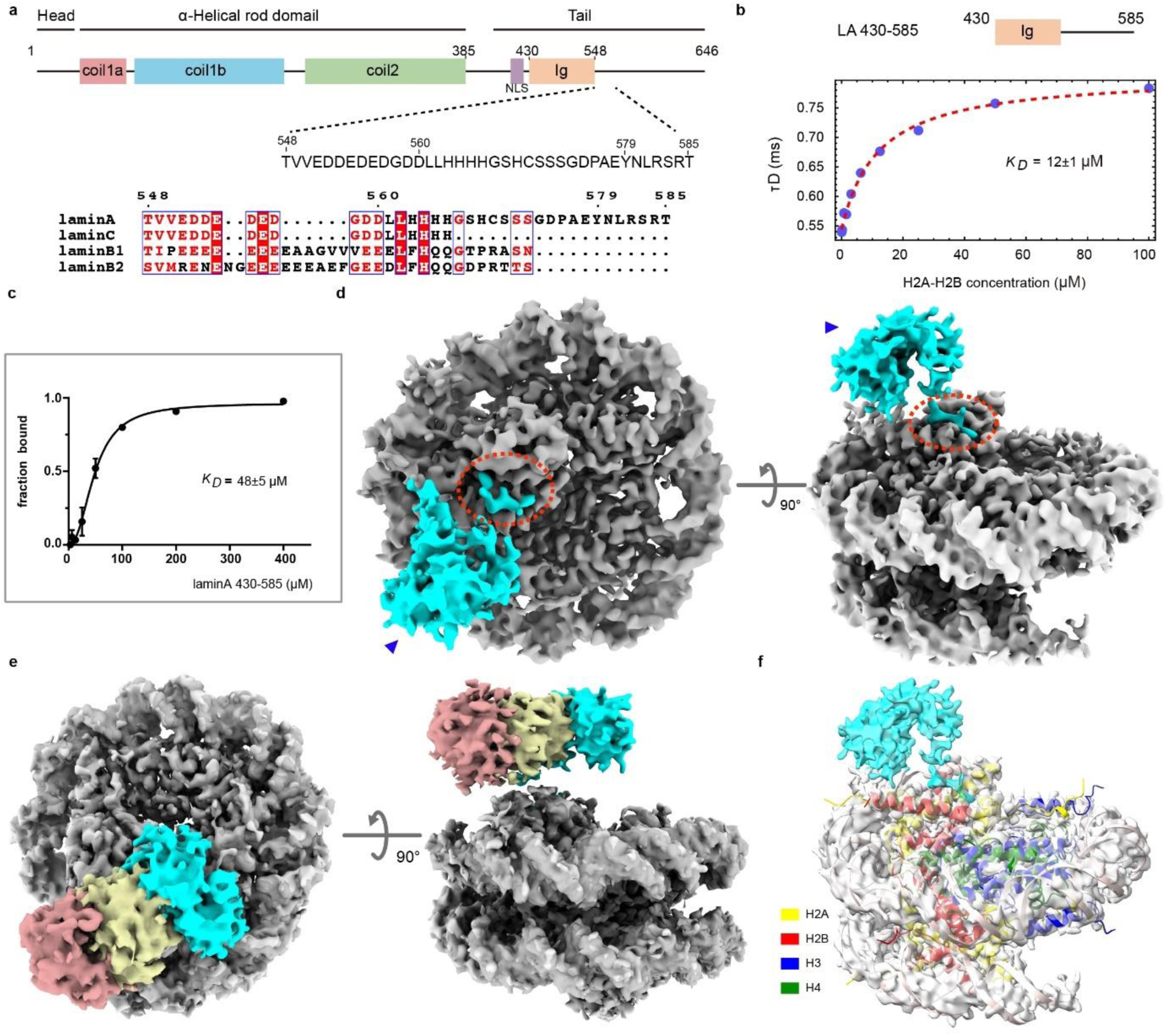
The tail domain of lamin A interacts with H2A-H2B heterodimers and nucleosomes. **a**. A schematic view of the lamin A protein shows the various protein domains and the position of the nuclear localisation sequence (NLS). The sequence of the lamin A tail domain, amino acids 548-585, is aligned to the other main lamin isoforms: lamin C, lamin B1, and lamin B2. The relatively conserved amino acids are coloured. **b.** The affinity of the truncated lamin A tail domain (LA 430-585) towards the H2A-H2B heterodimer was determined by FCS (Methods section, **Extended Data** Fig. 3c). **c.** The affinity between LA 430-585 and nucleosomes was quantified by an electrophoretic mobility shift assay (EMSA, **Extended Data** Fig. 4b). **d.** The structure of the LA 430-585 (cyan) – nucleosome (grey) complex was determined by cryo-EM. The density of the Ig-like domain (arrowhead) and additional lamin A tail density (red dashed circle) were identified. **e.** Variability in the position of the Ig-like domain was detected across different structural classes (reddish, yellow, and cyan), indicating flexibility between the stationary binding site (**d**. red dashed circle) and the Ig-like domain. **f.** The nucleosome structure (6ZHX) was fit into the EM density indicated that LA 430-585 interacts with the H2A-H2B heterodimer.

Next, we analysed the interactions between LA 430-585 and reconstituted nucleosomes^40^ (Extended Data Fig. 4a). Electrophoretic mobility shift assay (EMSA) showed direct binding between LA 430-585 and nucleosomes (Extended Data Fig. 4b), while no interactions were detected between nucleosomes and the truncated tail domain of lamin B1 amino acids 432-569 (LB1 432-569) (Extended Data Fig. 4c). We determined a K_D_ of 48 ± 5 µM for the binding of LA 430-585 to nucleosomes (Fig. 3c). This modest affinity suggests a dynamic interaction between lamin A and nucleosomes. Subsequently, using GraFix^41^, we purified the complex and conducted cryo-EM analysis. The structure of the LA 430-585 – nucleosome complex was resolved to ∼3.6 Å resolution using single particle cryo-EM (Fig. 3d, Extended Data Figure. 4 d-i). Two separate densities of the lamin A tail domain (Fig. 3d, cyan) were detected in addition to the nucleosome (Fig. 3d, grey). The characteristic density of the Ig-like domain (Fig. 3d, arrowhead) was found in multiple positions in the vicinity of the nucleosome (Fig. 3e, Extended Data Fig. 4f). The other density found attached to the nucleosome (Fig. 3d, red dashed circle) revealed a contact site with the H2A-H2B heterodimer of the nucleosome (Fig. 3f). We identified densities of a tyrosine and an arginine and assigned them to lamin A Y579 and R582. To validate these amino acid assignments, we utilised a shorter lamin A truncation LA 430-579 and analysed the structure of the LA 430-579 – nucleosome complex by cryo-EM (Extended Data Fig. 5a-f). As expected, the density map showed the canonical nucleosome structure without any additional density, implying that amino acids 579-585 are indispensable for the binding of lamin A to nucleosomes.

To further explore this interaction, we analysed the binding of a synthetic peptide comprised of amino acids 572-588 of lamin A (LA 572-588) to nucleosomes. Using cryo-EM structural analysis, we resolved the complex at 2.3 Å resolution (Fig. 4a, Extended Data Fig. 6a-e). Eight amino acids (AEYNLRSR) of the peptide were resolved at the nucleosome surface (Fig. 4b), while five amino acids (YNLRS) were found to interact with the acidic patch of the H2A-H2B heterodimer (Fig. 4c). The side chain of Y579 forms a hydrogen (H-) bond with H2A E56, stabilised by a hydrophobic interaction through H2A A60 and H2B V41. The main chain of N580 forms an H-bond with H2B H106 and E110. The C_β-γ_ of H2A E61, E64, the aromatic ring of H2AY57 and the side chains of H2A A60 and H2B V41 provide a hydrophobic surface for holding the side chain of L581. Most importantly, R582 acts as an arginine anchor to H2A E61, D90, and E92 through salt bridges. Finally, the side chain of S583 interacts with H2A E64 via a hydrogen bond. Remarkably, the YNLRS motif is unique to the lamin A tail domain (Fig. 3a) and highly conserved in evolution (Extended Data Fig. 3d).

**Fig. 4.**
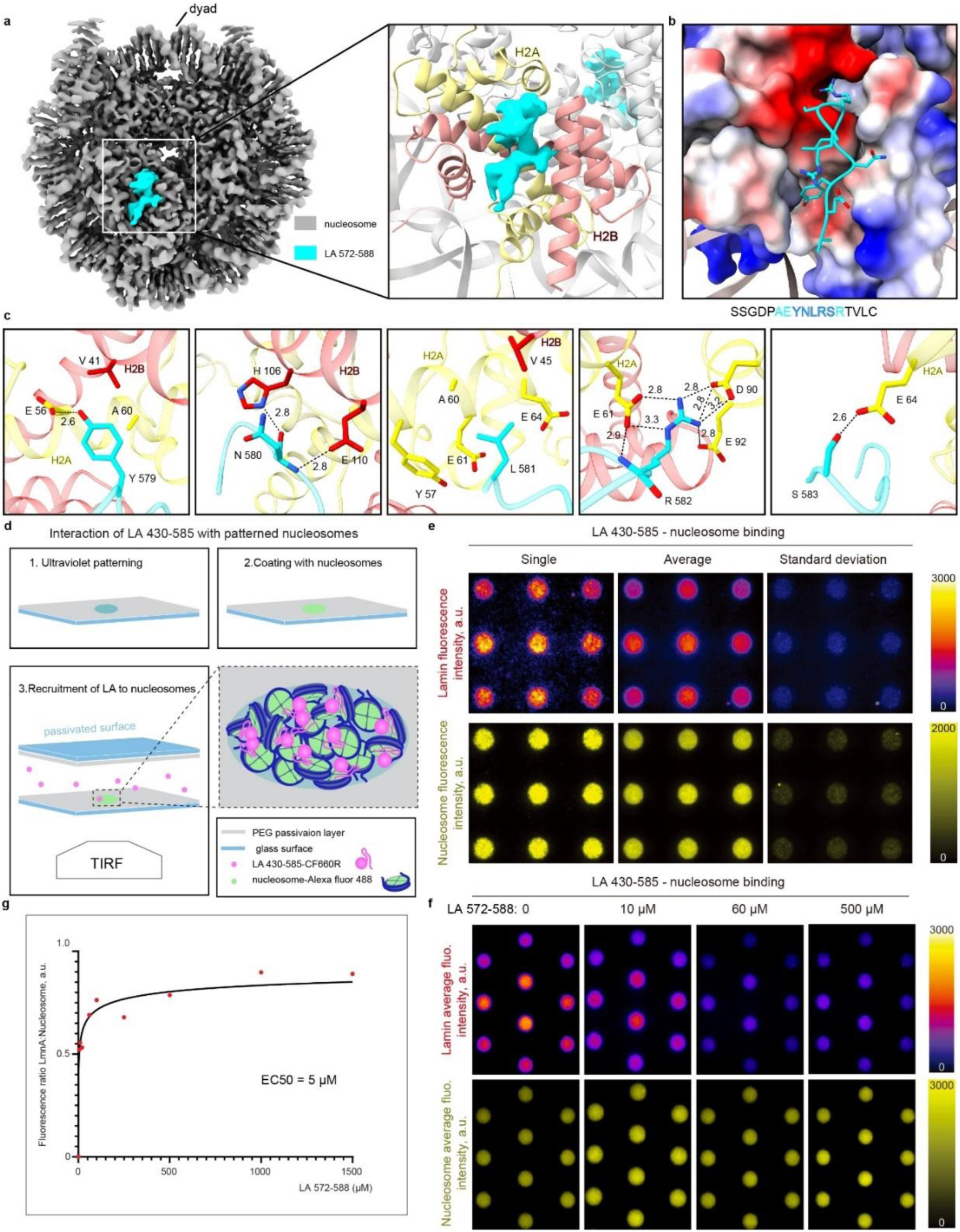
The amino acids 579-583 of lamin A mediate interaction with the nucleosome. **a**. Cryo-EM structural analysis of the peptide LA 572-588 (cyan) interacting with the nucleosome (grey) through the H2A-H2B heterodimer (yellow and reddish, respectively). **b**. 8 amino acids (AEYNLRSR) of the peptide LA 572-588 were identified in the negatively charged acidic patch of the nucleosome. **c.** Five amino acids YNLRS specifically interact with H2A-H2B within nucleosome. This interaction is stabilised by H-bonds, hydrophobic interactions, and salt bridges. **d**. A schematic view of the experimental design used for studying lamin A-nucleosome interactions by TIRF microscopy. A passivated glass coverslip was patterned using deep-ultraviolet illumination (1), then the patterned surface was coated with fluorescently labelled nucleosomes (2). The functionalised coverslip was mounted onto a pegylated glass slide to fabricate a chamber, where fluorescently labelled LA 430-585 polypeptide was injected at the onset of the binding reaction (3). **e**. The panels show the fluorescence intensity of the labelled LA 430-585 polypeptide (upper row) associated with patterned fluorescently labelled nucleosomes (bottom row) for representative single images, averaged images of 14 frames containing a total of 126 patterned spots, and related standard deviation images. **f.** The LA 572-588 peptide competes with the LA 430-585 polypeptide for nucleosome binding. The unlabelled peptide was added to the reaction mixture described in (**d**). The panels show the fluorescence intensity of the LA 430-585 polypeptide (upper row) associated with the patterned fluorescently labelled nucleosomes (bottom row) for average images of at least 24 frames. **g.** The fluorescence ratio of nucleosome-associated LA 430 – 585 against spotted nucleosomes was calculated (see Methods), and a curve was fitted with a four-parameter dose-response Hill-Langmuir equation (see Methods). The best fit of the overall dataset was obtained for a maximal ratio fixed to less than 1 and showed a half maximal effective concentration, EC50, of 5 µM. R2 was 0.78 for the overall dataset. **e, f** Fluorescence calibration bars are provided. All spots are 3 μm in diameter.

The nuclear lamina features a high concentration of lamin tail domains along each filament, which can potentially interact with dense nucleosome regions constituting the LADs. To mimic this scenario, we patterned surfaces with fluorescently labelled nucleosomes to create high concentration spots of nucleosomes. We utilised a fluorescently labelled LA 430-585 and exposed it to nucleosomes immobilised on patterned surfaces (Fig. 4d). Total Internal Reflection Fluorescence (TIRF) microscopy imaging showed that LA 430-585 binds specifically to the spotted nucleosomes (Fig. 4e), while LB1 432-569 does not, even at higher nucleosome concentrations (Extended Data Fig. 6f). Next, we demonstrated that the presence of the peptide LA 572-588, competed with LA 430-585 for nucleosome binding. This competition was evident from a decrease in fluorescence signals as a function of increasing peptide concentration (Fig. 4f). Quantification of these assays indicated an EC50 of 5 µM (Fig. 4g). While peptides commonly require higher concentrations for competition, this observation may hint at an additional low affinity of the tail domain to nucleosomes that accounts for the multiple positions of the Ig-like domain found in our structural analysis (Fig. 3e). Overall, these results show that the direct interaction between lamin A and nucleosome occurs via the tail domain. These interactions are dynamic, as expected for elements undergoing reorganisation during the cell cycle.

### A- and B-type lamins differently regulate chromatin architecture

To reveal the impact of lamins on genome organisation, we examined chromatin structure dynamics using genome-wide analyses, including 4f-SAMMY-seq^24, 42^, ChIP-seq^43^ and RNA-seq^44^. The 4f-SAMMY-seq method provides genome-wide solubility profiles, elucidating regions of open and closed chromatin and its compartmentalization into functional genomic regions (Extended Data Fig. 7a-d, Methods section). Positive values of the solubility profile coincide with euchromatin, while its negative values match heterochromatin, which contains LADs^42^, as evidenced by correlation analysis with ChIP-seq data of open and closed chromatin-associated modifications (Extended Data Fig. 7e, f).

By using 4f-SAMMY-seq we identified specific LmnaKO and LBDKO genomic regions exhibit significantly different solubility profiles compared to WT (Fig. 5 a-c, Methods section). Both lamin knockout strains presented with ∼ 10% genome-wide remodeling, with over half the change affecting heterochromatin organisation. Increased solubility of heterochromatin domains coincided with a decrease in H3K9me3-marked heterochromatin (Fig. 5a, b, Extended Data Fig. 7g, S3 DOWN). Exclusively in LBDKO cells, we observed spreading of the H3K9me3 signal to neighbouring DNA, suggesting that B-type lamins are necessary for the insulation of the H3K9me3-enriched heterochromatin regions (Fig. 5a, Extended Data Fig. 7g). The lamin knock-out cell lines showed little overlap of altered heterochromatic regions. On average, only ∼ 36% S3 DOWN and ∼ 40% S3 UP regions were shared between both knock outs (Extended Data Fig. 7h), accounting for 7% of the LmnaKO and LBDKO genomes having different solubility states (Fig. 5b). Notably, the X-chromosome exhibited an extensive loss of heterochromatin exclusively in LmnaKO cells (Extended Data Fig. 7i, S3 DOWN).

**Figure 5.**
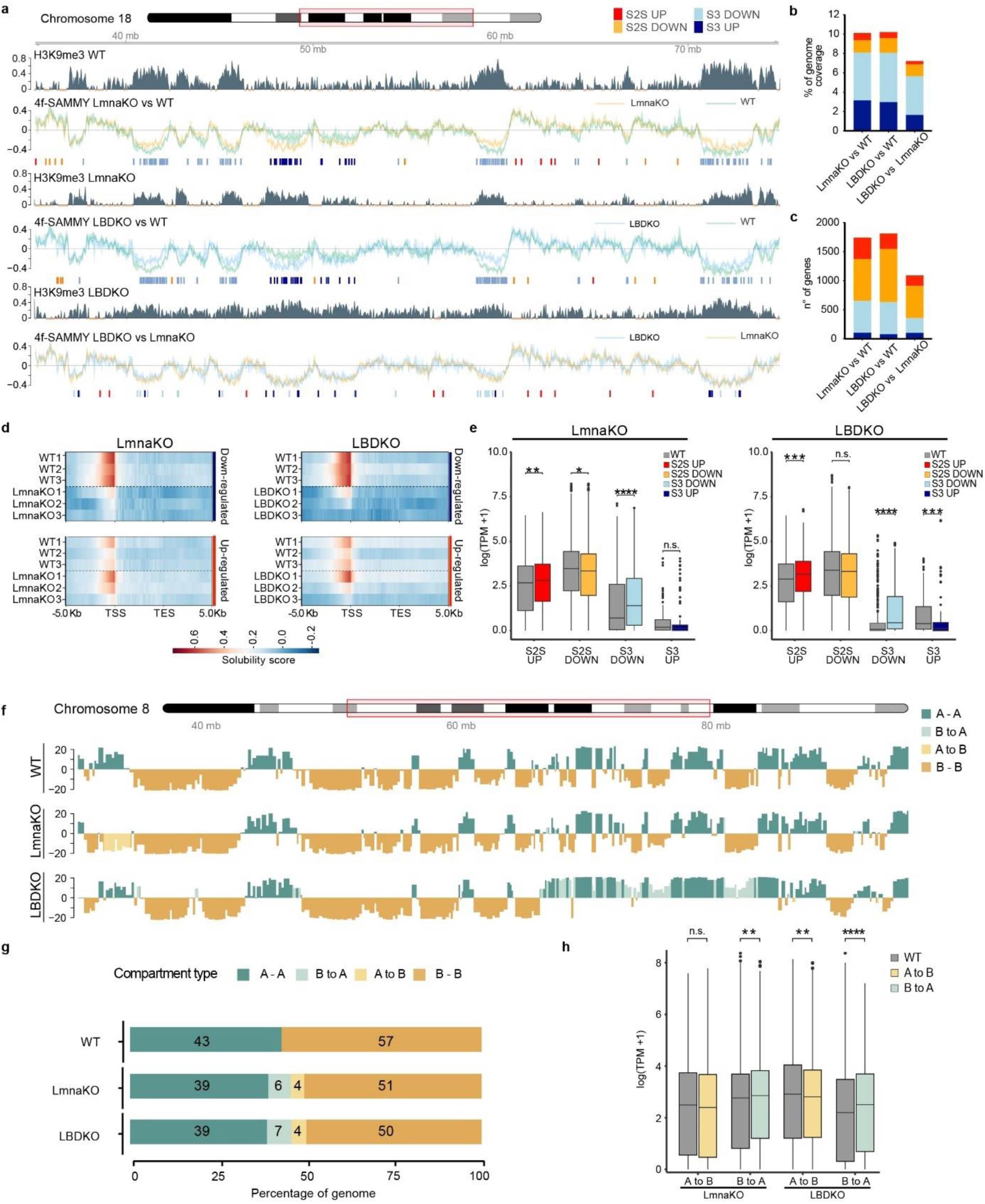
A- or B-type lamin specific heterochromatin remodelling. **a.** A representative genomic region of chromosome 18 (40 Mb on chr18: 35000000-75000000) showing tracks for chromatin solubility and H3K9me3 marks in the WT and lamin knockout cells. A representative H3K9me3 ChIP-seq is shown for each cell line (WT, LmnaKO, LBDKO). 4f-SAMMY-seq solubility profiles comparing the different cell lines are represented as the log of sequencing reads of the more soluble S2S over those of the less soluble S3. The line represents the mean of triplicates, and the standard deviation is shown as a shadow. Below each track pair, the respective significantly differentially soluble regions are indicated by coloured bars. **b.** The percentages of the genome affected by the removal of lamin genes are shown as a stacked bar plot, following the colour code described in **a. c.** Stacked bar plot showing the number of protein-coding genes with altered solubility (see Methods). **d.** Heat maps of solubility profiles of downregulated (above) and upregulated (below) Differentially Expressed Genes (DEGs) aligned to the transcription start and end sites (TSS and TES), shown as triplicates. Red and blue correspond to higher or lower accessibility, respectively. **e.** Box-plot distribution of log2 of transcripts per million (TPM + 1) in significantly differentially soluble regions in LmnaKO (left) and LBDKO (right). **f.** Chromatin compartment analysis using 4f-SAMMY-seq (see Methods). The representative first eigenvector on chromosome 8 (32,000,000-95,000,000) at 50 kb resolution is reported for WT, LmnaKO, and LBDKO cells. Regions with concordant (A-A, and B-B) or discordant compartment (B to A, and A to B) classification in LmnaKO and LBDKO compared to WT are marked. **g.** A stacked bar plot showing average percentages of compartment switching across genomes in comparison to WT. **h.** Box-plot distribution of the log2 of transcripts per million (TPM + 1) calculated for genes of discordant compartment with respect to WT. In **d, h**, the box lower and upper edges are the first and third quartiles, and the horizontal bar is the median. Whiskers extend up to 1.5 times the interquartile range (IQR) from the edges. Data points outside the range are outliers and are represented by dots. The statistical significance of the data has been assessed using the Wilcoxon rank sum test labelled as p-value<0.0001(****), p-value<0,001(***), p-value<0.01(**), p-value<0.05 (*), and ns p-value>0.05.

To correlate these findings with transcriptional regulation, we performed RNA-seq. The differential expression analysis showed that ∼ 40% of upregulated and ∼ 30% of downregulated genes were commonly regulated by A-type and B-type lamin knockouts (Extended Data Fig. 8a, b). Gene ontology (GO) analysis of Differentially Expressed Genes (DEGs) detected terms, indicating aberrant regulation of spatial-temporal gene expressions in the absence of lamin proteins (Extended Data Fig. 8c, Extended Data Table 3). By integrating the RNA-seq with the solubility analysis, we found that downregulated genes in lamin knockout cells exhibit lower chromatin solubility (Fig. 5d). However, we did not find a clear increase in solubility for upregulated genes, suggesting that transcription is also compatible with a low solubility state. To further understand the link between solubility and transcription, we quantitatively analysed the transcription of all genes localised in significantly differentially soluble regions (Fig. 5e). Transcription quantitatively changes in accordance with chromatin solubility variations, supporting the view that functional genomic changes are mediated by lamin-induced heterochromatin reorganisation.

Based on the biochemical properties of chromatin, we built 3D compartmentalisation maps (see Methods section), as seen by HiC^45^. Chromatin was segregated into active ‘A’ and inactive ‘B’ compartments for WT and lamin knockout cell lines (Fig. 5f, Extended Data Fig. 8d). Interestingly, although knockout of A- and B-type lamins showed a similar degree of compartmentalisation shifts (Fig. 5g), half of the affected regions were exclusively influenced by either A-type or B-type lamins (Extended Data Fig. 8e). This was reflected in the transcriptional changes within the A and B compartments (Fig. 5h). In line with previous solubility analysis (Fig. 5e), transcription associated with heterochromatin loss is more significant. GO analyses revealed that genes that underwent compartment shift upon knockout of lamins were likely related to distinct biological functions. These results suggest that A-type and B-type lamins epigenetically regulate common and lamin-specific domains of the genomic compartments, implying that they may influence distinct epigenetic regulations (Extended Data Fig. 8f, Extended Data Table 4).

## Discussion

Lamin-chromatin interactions impact nuclear functions as well as gene expression^46^. Using state-of-the-art *in situ* and *in vitro* approaches, we shed light a on how lamin isoforms interact with nucleosomes and can modulate their distribution at the nuclear lamina and around the nuclear pores, impacting heterochromatin compartmentalization and transcription regulation. A model summarising our findings is shown in Figure 6. Our biochemical analysis has indicated that the tail domain of lamin A can directly bind to purified H2A-H2B heterodimers, as well as assembled nucleosomes. Lamin A interacts with nucleosomes through the acid patch of H2A- H2B, which accounts for its property to bind to purified H2A-H2B heterodimers. While the interactions of lamins with purified histones have been detected in vitro^47^, nucleosome-lamina interactions are more physiologically relevant. Here, we identified the motif YNLRS in lamin A specifically recognise the nucleosome (Fig. 6, upper left). This interaction is unique to lamin A and appears to be evolutionarily conserved, as other lamin isoforms lack the YNLRS sequence motif. Hence, this study suggests a functional distinction between lamin A and other isoforms. Although lamin A tail-nucleosome interactions exhibit a relatively modest affinity, the high concentration of lamin A tail domains at NL and the dense nucleosome organisation in LADs allow this interaction. Consequently, this leads to reduced dynamics of lamin A compared to lamin C, as shown previously^48^. Indeed, the removal of a fragment of lamin A that contains LA 572-588 increases its mobility at the NL^49^ .

**Figure 6.**
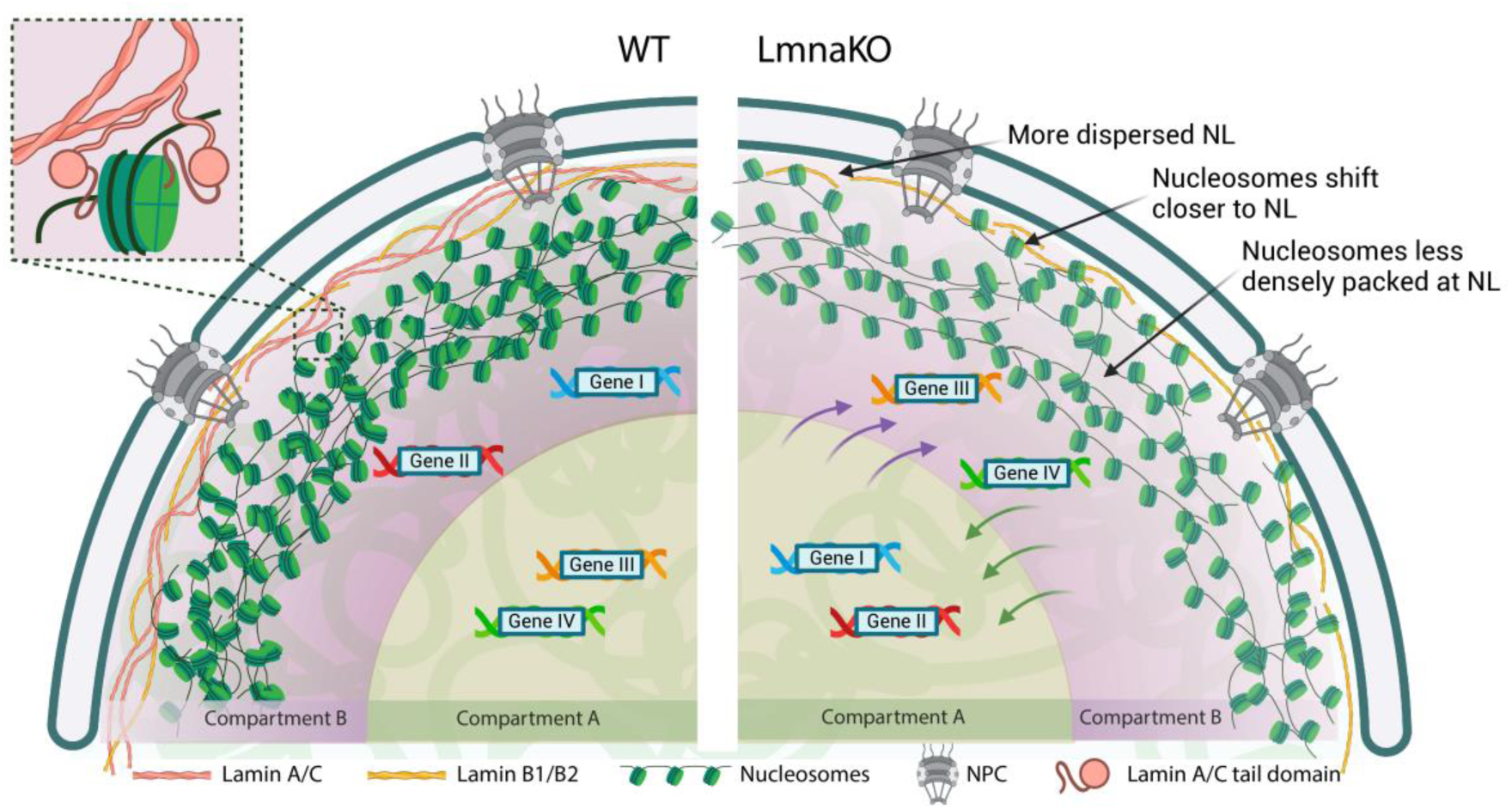
Schematic model of the NE in WT and LmnaKO MEFs. The model shows lamins and chromatin organisation underneath the NE in WT and LmnaKO MEF cells. The chromatin (nucleosomes in green) forms a dense structure at the NE by association with the NL, consisting of Lamin A/C (red) and Lamin B1/B2 (yellow) in WT cells. Lamin A can interact with the nucleosomes (upper left), via its tail domain. In the absence of A-type lamins (LmnaKO), chromatin remodelling affected the concentration of nucleosomes at the NE, which are less densely packed and shifted closer to the more dispersed nuclear lamina. These structural effects lead to major nuclear reorganisation, including a large number of genes moving from transcriptionally active compartment A to inactive compartment B, and vice versa.

The lamin A tail domain is composed of two intrinsically disordered amino-acid stretches, 45 and 98 amino acids long, that are separated by a well-structured Ig-like domain (Fig. 3a). These long, intrinsically disordered regions would permit the lamin A tail domain to extend up to ∼50nm in length. Thus, we estimate that lamin A may directly interact with nucleosomes located up to 30-35nm from the NL, due to the long amino acid stretch of the tail up to the binding site. This would allow interactions of the tail domain with multiple components i.e., while interacting with the nucleosome, the Ig-like domain can potentially interact with other components such as BAF. Importantly, lamins can also interact with chromatin via their binding proteins, e.g., Emerin, LEMD2 and LBR^50^.

The organisation of the chromatin at the NE allows cells to regulate gene expression and other essential nuclear functions^51^. Here, we revealed the alteration in spatial nucleosome concentration at the NE and as a consequence of the knockout of specific types of lamins in MEFs. Chromatin at the NL, primarily consisting of LADs, is enriched with heterochromatin histone methylation marks and only hosts a small fraction of expressed genes^52^. However, our data revealed a variation in the concentration of nucleosomes at the NE, which may correlate with more open chromatin, typically associated with expressed genes.

Nuclear lamins and chromatin synergistically form a mechanical functional unit to overcome cellular and environmental stress^53^. This synergy relies on tight interactions between these two structures. Cells that express different isoforms of lamins show variations in the lamin meshwork. Notably, we found a reduced concentration of nucleosomes, i.e., chromatin, at the nuclear lamina of MEF cells that express solely B-type lamins. This coincides with previous observations indicating that A-type lamins are mostly expressed in differentiated cells, where the interactions of tightly packed heterochromatin with the NL are enhanced^54^. Consistently, only modest changes in nucleosome concentration were detected upon removal of B-type lamins, suggesting that A-type lamins exert a prominent influence on chromatin organisation at the nuclear envelope (Fig. 6).

Genome-wide chromatin analysis revealed that A- and B-type lamins have a selective role in the functional organisation of the genome, predominantly by controlling heterochromatin organisation (Fig. 5). In agreement with our cryo-ET analysis (Fig. 1), it was shown that reduction in A-type lamin expression resulting in local changes of LADs, reduction of H3K9me3 levels^55^. Here, we found that A-type lamins impact heterochromatin solubility and impair transcription from these regions. Interestingly, the absence of B-type lamins leads to a loss in H3K9me3 domain boundaries. Overall, A- and B-type lamins displayed distinct patterns of transcriptional regulation, suggesting their involvement in specific epigenetic pathways, underlying the importance of native lamin expression in retaining functional nuclei. Our analysis indicated that loss of lamin A/C dependent heterochromatin compartments leads to epigenetic alteration of genes that participate in histone regulation. These results may explain the decrease in nucleosome concentrations found at the nuclear envelope. Moreover, it suggests a substantial alteration of chromosome organisation in agreement with single MEF cell fluorescence analysis^56^ and previous genome-wide analysis^57^.

In summary, we used *in vitro* and *in situ* structural analysis to reveal the molecular basis of lamin-chromatin interactions in conjunction with genome-wide analysis. Our findings demonstrate the lamin-dependent organisation of the 3D genome and its impact on gene expression variability. A- and B-type lamins exhibit differential regulatory roles in chromatin architecture, with lamin A distinguished by its direct interaction capabilities with nucleosomes. Therefore, retaining the native levels of each lamin isoform is important for the optimal function of cells.

## Author contributions

B.W. prepared lamin truncations, nucleosome reconstitution, cryo-EM data acquisition and analysis with the help of Q.L. R. K-T. prepared, acquired and analysed the cryo-FIB cryo-ET including the sub-tomogram averaging of nucleosomes with the help of M.E., R.B.P. conducted the TIRF experiments and analysed the data. V.R. studied chromatin solubility and RNA analysis with the help of E.D.P.S. who conducted the required bioinformatics. L.P. and B.S. conducted and analysed the FCS affinity experiments. C.L. conceived the research and secured funding. O. M. conceived the research, supervised the project, secured funding and wrote the manuscript with contributions from all authors.

## Data availability

The cellular tomograms and sub-tomogram averaging structures were deposited in the EMDB: EMD-19827, EMD-19828, EMD-19829, EMD-19824. The density map of lamin A 430-585 with nucleosome was deposited in EMDB: EMD-50291. The complex structure of lamin A 572-588 with nucleosome was deposited in the EMDB: EMD-50114 and PDB: 9F0O. The high-throughput sequencing data generated for this study are available in the NCBI GEO database with following accession numbers ‘GSE268922, GSE268923, GSE268924’.

## Acknowledgement

This work was funded by grants from the Swiss National Science Foundation (SNSF 391 310030_207453) to O.M. and (SNSF 310030_197776) to B.S., AFM (grant #24306); FRRB (grant # 3444218) and MIUR (PRIN #2022-4RFLLA) to C.L. We thank the Center for Microscopy and Image Analysis (ZMB) at the University of Zurich and Vittoria Moretti, Daniele Marchelli of the sequencing facilities of Fondazione IRCCS Ca’ Granda-Ospedale Maggiore Policlinico, in Milan. We thank Alice Thurston for critical reading of the manuscript.

## Methods

### Cells and cell culture

Wild-type, Lmna^-/-^, and Lmnb1^-/-^+Lmnb2^-/-^ MEFs were grown at 37 °C in a humidified incubator with 5% CO2 in high glucose DMEM (Sigma-Aldrich, D5671) supplemented with 10% FCS (v/v), 2 mM L-glutamine (Sigma-Aldrich, G7513) and 1% penicillin-streptomycin (Sigma-Aldrich, P0781). Cells were seeded onto glow discharged cryo-EM grids (Quantifoil 2/1, 200 mesh, Au) overnight. Grids were washed in 1xPBS (Fisher bioreagents, BP399-1) before manual plunge freezing into liquid ethane cooled by liquid nitrogen. Grids were stored in liquid nitrogen before further use.

### Cryo-FIB milling

Grids were clipped into CryoFIB AutoGrids (Thermo Fischer Scientific, USA) before being transferred to the FIB-SEM loading station. The grids were mounted onto a custom-built grid holder before being sputter coated with 8 nm Pt/C in a Leica BAF060 system cooled to -160 °C and directly inserted into the Zeiss-Auriga 40 Crossbeam FIB-SEM with a stage temperature of -150 °C. Before milling, grids were covered with 2x5 s flashes of organometallic platinum with the internal gas injection system. Milling was performed at a stage angle of 18 ° and a FIB beam current from 240 to 20 pA at a constant voltage of 30 kV. The process was controlled with the Nano Patterning and Visualization Engine software (Zeiss) and progress was observed with the SEM. Grids with finished lamellae with a thickness of 100–200 nm were stored in liquid nitrogen until further usage.

### Cryo-electron tomography

Grids with FIB-milled lamellae were transferred to a Titan Krios electron microscope (Thermo Fischer Scientific, USA) equipped with a Gatan K2 summit direct electron detector and a Gatan energy filter operated at 300 kV in zero-loss mode. Tomograms of the nuclear envelope were acquired with a magnification of 64000x (0.21 nm pixel size) with -4 µm defocus and a dose symmetric tilt scheme^58^ from ±60 ° in 3 ° increments and a total dose of 160 e^-^/Å^2^ in SerialEM^59^.

Drift-corrected tilt series were aligned and CTF corrected with IMOD^60^ using either platinum depositions as fiducials or the IMOD patch tracking for alignment. For subsequent lamin and nucleosome detection, tomograms were reconstructed with WBP in bin 4 with a SIRT-like filter applied.

### Cryo-ET image processing and data analysis

Nucleosomes were detected using a crYOLO^61^ model trained on a data subset of 10 tomograms and applied on the full dataset from all three cell lines. This resulted in 130000 nucleosome coordinates, which were imported into MATLAB (MathWorks, USA) using the TOM toolbox^62^ where subtomograms of 16x16x8 pixels were projected in z to generate 2D particles. An initial 2D classification was performed with RELION 4.0.1^33, 63^ to remove false positive nucleosome detections. The resulting coordinate set of 101780 nucleosomes was then processed using the subtomogram workflow of RELION 4.0.1^33, 63^. Pseudo-subtomograms were subjected to a 3D refinement using PBD 7XD1^64^ filtered to 30 Å as an initial reference. The aligned coordinates from the 3D-refined structure were imported to MATLAB (MathWorks, USA) and used for all further measurements.

Lamins were manually segmented using IMOD’s^60^ 3dmod software. To be able to properly follow the filaments, the tomograms were rotated using the slicer viewer to bring the lamina into the viewing plane. Resulting filament coordinates were imported into MATLAB (MathWorks, USA) and resampled into equidistant coordinates with a 3.5 nm distance using the interparc function^65^. This resulted in 44880 lamin coordinates across the three cell lines. Additionally, in the tomograms containing one or more NPCs, the centres of the NPCs were defined as coordinates for further measurements.

All distance and neighbourhood measurements were performed in MATLAB (MathWorks, USA). For all measurements, we excluded nucleosomes that were further away from the nuclear lamina than 100 nm because of the different field of views between tomograms and to focus on the lamin-peripheral nucleosomes. All neighbourhood measurements were done by measuring the number of neighbours in a sphere with a 24 nm radius.

Graphs were drawn with Origin 2018 (OriginLab, USA) and 3D visualization was performed with ChimeraX^66^. Additionally, the Artiax^67^ plugin was used for the NPCs.

### Histone protein expression, purification and heterodimer assembly

*Xenopus laevis* histones, H2A, H2B, H3, and H4, were expressed and purified as previously described^40^. H2A and H2B or H3 and H4 were mixed separately at equal molar ratios, followed by dialysis using refolding buffer (2M NaCl, 20mM Tris-HCl pH 8.0, 1 mM DTT). Insoluble components were pelleted by high-speed centrifugation. Soluble H2A-H2B heterodimers and H3-H4 tetramers were purified using size exclusion chromatography (SEC).

### DNA preparation, nucleosome assembly and purification

The 197 bp DNA fragment was designed based on the 147 bp 601 wisdom nucleosomal DNA ^68^and cloned into the PUC57 plasmid. Large-scale PCR amplification was performed, followed by purification using the phenol-chloroform DNA extraction method. DNA was then precipitated in 70% ethanol and resuspended in 20 mM Tris-HCl pH 8.0, 0.5 mM EDTA buffer (TE buffer) for subsequent nucleosome reconstitution.

*Xenopus laevis* histones were assembled into nucleosomes as previously described^40^. Following the removal of dissolved components by centrifugation, the supernatant was concentrated and subjected to further purification using a Superdex^TM^ 200 Increase 10/300 GL column (cytiva). The fractions obtained were analysed by SDS-PAGE.

### Fluorescence labelling of nucleosome with Alexa Fluor™ 488

Nucleosomes containing a cysteine substitution at H4K20 (H4K20C) were reconstituted and purified using established protocols. Then the H4K20C nucleosome was dialysed into PBS buffer, followed by two rounds of dialysis. The labelling process involved incubating the H4K20C nucleosomes with Alexa Fluor™ 488 C_5_ Maleimide (Thermo Fisher Scientific, A10254) dye at a 5:1 molar ratio overnight at 4 °C. To quench the labelling reaction, DTT was added to the sample at a final concentration of 10 mM. Finally, free dye was separated from the labelled nucleosomes using glycerol gradient ultracentrifugation, using a 10% to 30% glycerol gradient.

### Constructs of lamin tail domains and protein purification

DNA sequences encoding for lamin A 394-646 (LA 394-646), 394-548 (LA 394-548), 430-585 (LA 430-585), 430-579 (LA 430-578), 430-560 (LA 430-560), laminB1 432-569 (LB1 432-569) were cloned into pET28 vector. Additionally, DNA sequences for LA 430-585 and LA 430-579, intended for structural analysis, were inserted into the 6xHis-SUMO plasmid. For FCS experiments and TIRF microscopy, the LA 430-585 C570A mutation was prepared in the 6xHis-SUMO plasmid to allow specific labelling of CF660R SE dye at position C522. Lamin B1 432-569 (LB1 432-569) was also cloned into the same plasmid.

The above constructs were transformed into the BL21(DE3) bacterial strain and cultured in TB medium at 37 ℃. Protein expression was induced by adding 0.5 mM IPTG, followed by 16 h culturing at 18 ℃. Harvested bacteria were lysed into 500 mM NaCl, 20 mM Tris-HCl pH 8.0, 25 mM imidazole buffer. Bacteria were lysed by sonication followed by centrifugation at 20,000 g for 1 h. The supernatants were applied to Ni-NTA affinity chromatography. Targeted proteins were eluted by a linear gradient with elution buffer (500 mM NaCl, 20 mM Tris-HCl pH 8.0, 500 mM imidazole). The His-SUMO-tagged proteins were subjected to ULP1 protease incubation, followed by dialysis into the imidazole free lysis buffer for 3 h. Next, the samples were purified using Ni-NTA affinity chromatography, allowing the target proteins to be collected and concentrated to the desired concentration. Finally, proteins were further purified by size exclusion chromatography using a HiLoad^TM^ 16/600 Superdex^TM^ 75 pg (cytiva)., The buffer contained 150 mM NaCl, 20 mM HEPES pH 7.5, 2 mM DTT. The samples were accessed by SDS-PAGE, concentrated, and stored at -80 ℃.

### Histone H2A-H2B-laminA tail binding assay

Purified LA 394-646, LA 394-548, LA 430-585, LA 430-579, LA 430-560, along with a His-GST (negative control), were immobilised on the Ni-NTA resin using a buffer containing 150 mM NaCl, 20 mM HEPES pH 7.5, 25 mM imidazole, 0.01% tween. Rigorous washing steps were performed prior the introduction of equimolar amounts of the H2A-H2B heterodimer to each lamin truncation. The samples were incubated for 1 h followed by extensive washing steps. Subsequently, the resin was boiled and analysed by SDS-PAGE. Equal protein amounts were used for each truncation to ensure an accurate comparison.

### Glass surface passivation and deep-ultraviolet (UV)-mediated micropatterning

Prior to passivation, slides and coverslips (CVs) were drastically cleansed by successive chemical treatments: 2 h in 2% Hellmanex III; rinsing in ultrapure water; 30 min in acetone; 30 min in 96% ethanol; rinsing in ultrapure water. Slides and CVs were dried using a filtered nitrogen gas flow and oxidised with oxygen plasma (3 min, 30 Watt, Femto low-pressure plasma system Type A, Diener electronic GmbH, Germany), just before an overnight incubation in a solution containing tri-ethoxy-silane-PEG (5 kDa, PLS-2011, Creative PEGWorks, USA) 1 mg/ml in 96% ethanol and 0.02% of HCl. mPEG-silane passivated slides and CVs were stored in a clean container and used within a week.

To direct the binding of nucleosomes to predefined positions on glass CVs, we used the micropatterning strategy^69^ by printing adhesive patterns on a protein-repellent surface. mPEG-silane passivated CVs were exposed to short-wavelength UV radiation (184.9 nm and 253.7 nm, Jelight, USA) for 2 min through 24 x 24 transparent micropatterns printed on a photomask (Compugraphics, Germany), and immediately mounted onto a PEGylated slide using a double-sided tape (3M electronics). Then, immediately, the fabricated flow chamber was incubated for 5 to 10 min with 0.2 µM of 25% Alexa fluor 488 labelled nucleosomes, then saturated with 0.8% BSA in a wash buffer containing 20 mM HEPES pH 7.5 and 100 mM NaCl, and finally washed in the wash buffer supplemented with 0.08% BSA. Likewise, free nucleosomes were washed away, and any uncoated position was likely saturated with BSA.

### TIRF Microscopy imaging

Reconstitution assays were performed using freshly functionalised chambers and the Lamin A/nucleosome reaction buffer, containing 20 mM HEPES pH 7.5, 100 mM NaCl, 0.3% BSA, 0.03% TWEEN 20, 60 mM β-mercaptoethanol, 0.6% methylcellulose, and lamin polypeptides. The final CF660R fluorescently labelled LA 430 – 585 truncation polypeptide concentrations were 1 µM (Fig. 4e). Alternatively, nucleosome-coated patterns were also tested for their binding capacity to the CF660R fluorescently labelled LB1 432 – 569 truncation polypeptides at a final concentration of 1 µM (Extended Fig. 6f). The reaction medium was also supplemented with a variable concentration of unlabelled lamin A-derived peptide 572 – 588. This reaction medium was injected into a passivated flow cell at the onset of the reaction, and imaging was performed after 1 h, at steady state. Specifically for Figure 4, the amount of nucleosome-associated lamin A truncation polypeptide 430 – 585 was calculated from 442, 815, 560, 369, 731, 653, 675, 665, 616, 298 spots for 0, 5, 10, 20, 60, 100, 250, 500, 1000, 1500 µM peptide 572 – 588, respectively, with a SEM of 0.007, 0.003, 0.010, 0.006, 0.002, 0.003, 0.002, 0.002, 0.002, and 0.004, respectively. TIRF images were acquired using a Widefield/TIRF – Leica SR GSD 3D microscope, consisting of an inverted widefield microscope (Leica DMI6000B / AM TIRF MC) equipped with a 160x objective (HCX PL APO for GSD/TIRF, NA 1.43), a Leica SuMo Stage, a PIFOC piezo nanofocusing system (Physik Instrumente, Germany) to minimise the drift for an accurate imaging, and combined with an Andor iXon Ultra 897 EMCCD camera (Andor, Oxford Instruments). Fluorescent proteins were excited using 3 solid-state diode lasers, 488 nm (300 mW), 532 nm (500 mW), 642 nm (500 mW). Laser power was set to 5% for Alexa-labelled proteins, and dyes were excited for 50 ms. Image acquisition was performed with 25 degrees-equilibrated samples and microscope stage. The microscope and devices were driven by Leica LAS X software (Leica Microsystems, GmbH, Germany).

### Image processing and data analysis of fluorescence images

Steady-state images taken after 1 h reaction was processed with Fiji software (NIH). 46 to 88 or 54 steady-state images were taken for LA 430-585 polypeptide +/- LA 572–588 peptide or LB1 432-569 polypeptide, respectively. To determine the ratio of nucleosome-bound lamin polypeptide to the total patterned nucleosomes, macros written in Fiji allowed to first subtract the background for each individual image and quantify the mean fluorescence intensity of lamin polypeptides and related nucleosomes immobilised on patterned spots in each individual image. The fluorescence ratio was then calculated, and data (Fig. 4f) was fitted with a dose–response curve using GraphPad (“[Agonist] vs. response, Variable slope”). The equation was:

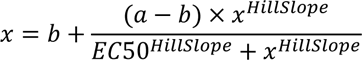

where a and b are plateaus in the units of the Y-axis, HillSlope describes the steepness of the curve, EC50 is the concentration that gives a response halfway between a and b. Three parameters, EC50, HillSlope, and b, were unconstrained. The best-fit values for the four parameters, plateaus, Hill slope, and EC50 were calculated from the overall dataset, with the value of the saturation plateau set to less than 1.

### Fluorescence correlation spectroscopy (FCS)

The CF660R SE (Sigma-Aldrich, SCJ4600053) labelled LA 430-585 C570A and LB1 432- 569 were subjected to SEC to remove free dye. FCS measurements were conducted at protein concentrations of 500 pM in a buffer composed of 20 mM HEPES pH 8 and 150 mM NaCl, supplemented with 0.01% Tween20 (to reduce surface adhesion) and 2 mM of both ascorbic acid and methyl viologen as photoprotectants. The samples were measured in µ-Slide chambers (ibidi) on a custom-built confocal single-molecule fluorescence instrument equipped with a UplanApo 60x/1.20 W objective (Olympus) and a red 640-nm diode laser (LDH-D-C-640, PicoQuant) used in continuous-wave mode at a power of 50 µW. Fluorescence photons were separated from scattered light by a triple-band mirror (zt405/530/630rpc, Chroma) and then passed through a 100-μm pinhole. CF660R photons were selected by a dichroic mirror (T635LPXR, Chroma) and an LP647RU long-pass filter (Chroma) and detected by a SPCM-AQRH-14 single-photon avalanche diode detector (Excelitas).

Data analysis was performed using Mathematica (Wolfram Research) with the custom add-on Fretica (available at https://schuler.bioc.uzh.ch/programs/). Correlation curves, *G*(*τ*), were fitted with a model that includes an amplitude, 1/*N*, and a diffusion time, *τ_D_*, for each measurement, and triplet dynamics (with a triplet amplitude, *c_T_*, and a triplet time, *τ_T_*) that is common for all points of a titration:

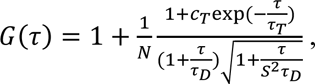

where *s* = 6 is the ratio of the axial to lateral radii of the confocal observation volume. The extracted diffusion times, *τ_D_*, were plotted against H2A-H2B concentration, and this curve was fitted with a one-to-one binding model between LA 430-585 C570A and the H2A-H2B dimer. The fit parameters were the equilibrium dissociation constant, *K*_D_, the diffusion time of unbound LA 430-585 C570A and the diffusion time of H2A-H2B-bound LA 430-585 C570A.

### Electrophoretic mobility shift assay (EMSA)of nucleosomes-lamin complexes

EMSA was conducted at 4 ℃ in a buffer containing 150 mM NaCl, 20 mM HEPES pH 7.5, 1 mM DTT. Alexa 488-labelled nucleosomes were utilised to assess binding to LA 430-585 and LB1 432-569. A concentration of 0.05 µM of labelled nucleosomes was incubated with progressively doubling amounts of either LA 430-585 or LB1 432-569. The concentration range of LA 430-585 or LB1 432-569 spanned from 3.1 µM to 400 µM. Following 1 h incubation, the samples were loaded onto a 6% native PAGE gel and imaged for 488 nm fluorescence using Fusion Fx Spectra (Vilber). The decreased quantity of nucleosomes was calculated by ImageJ and the curves were fitted by GraphPad Prism 10.

### Cryo-EM grid preparation and data collection

An excess of LA 430-585 was mixed with purified nucleosomes and incubated on ice for 1 h. Grafix method^41^ was applied to prevent aggregation and stabilise the lamin-nucleosome complex. The glycerol gradient ranged from 10% to 30% while the gradient of glutaraldehyde ranged from 0 to 1% in a buffer containing 100 mM NaCl 20mM, 20 mM HEPES pH 7.5. A total of 200 µl of mixed the complex was loaded and centrifuged at 215,600 g for 16 h at 4 ℃. The gradient was manually fractionated using 200µl per fraction and Tris-HCl pH 8.0 was added into each aliquot to quench the excessive crosslinking. 10 µl was uploaded into 6% native PAGE, to identify the position of the complex. Fractions containing the complex were selected, and the buffer was changed into 100 mM NaCl and 20 mM HEPES, pH 7.5. As a control, LA 430-579 was prepared in a similar manner. For peptide bound nucleosome complex, 1 mM of the 17-amino-acid peptide (SSGDPAEYNLRSRTVLC, PEPTIDE 2.0 Inc.) was mixed with 1 µM nucleosome in 100 mM NaCl, 20mM HEPES, pH 7.5 buffer.

Vitrified grids were prepared by applying 3 µl of 1 uM complex sample to a freshly glow-discharged holey carbon grids (Quantifoil R1.2/1.3 Au 200 mesh). The sample was blotted for 3-5 s at 4 ℃ with 100% humidity and plunged-frozen in liquid nitrogen cooled ethane (FEI ^TM^, Vitrobot). Electron micrographs were acquired using a 300kV Titan Krios G3 (Thermo Fisher Scientific) equipped with a BioQuantom and K3 direct electron detector (Gatan) in super-resolution mode. All images were recorded by EPU (Thermo Fisher Scientific) with a pixel size of 0.65 Å and a defocus range from 0.8 µM to 2.8 µM with a total exposure of ∼70 e^−^/Å^2^.

### Cryo-EM data processing

Datasets were processed by CryoSPARC v4.4.1^70^ following the workflow in Extended Data Fig. 4,5,6. Initially, all movies were subjected to live processing. Patch motion correction and patch CTF estimation were applied to the movies. High-quality micrographs were sorted out based on relative ice thickness and CTF resolution. Subsequently, particles were picked using automated template picking and extracted with a box size of 360 pixels, which were then downsampled to 180 pixels for following 2D classification.

For the LA 430-585/LA 430-579 bound to nucleosome complex, particles selected after several rounds of 2D classification were utilised for generating three or two ab initio classes, followed by heterogenous refinement. In the case of the peptide-nucleosome complex, particles were used for generating two ab initio classes. Promising classes were subjected to focused 3D classification based on the Ig-like domain density.

Classes of interest were further processes through homogeneous refinement, non-uniform refinement, reference-based motion correction, another round of non-uniform refinement, local CTF refinement, and non-uniform refinement. The quality of the map was validated by FSC validation and local resolution analysis.

For the peptide-nucleosome structure, the initial model was obtained by fitting a well-solved nucleosome model from 6ZHX^71^ into the final map using ChimeraX^66^. The model of the nucleosome was refined into cryo-EM density using Phenix. The model of peptide was manually built in Coot^72^ and refined in PHENIX^73^ iteratively. Visualization of all cryo-EM maps and figure preparation were done by Chimera and ChimeraX^66^.

### Cell-cycle assay

2x10^5^ cells were collected, rinsed with PBS, and then fixed in 80% ice-cold ethanol at 4 °C. The fixed cells were washed twice with ice-cold 1xPBS, followed by incubation in 1xPBS containing 25 mg/ml propidium iodide (PI, Sigma, P 4170) and 0.5 mg/ml RNase A (Invitrogen, EN0531) for 1 h at 4 °C. Following staining, the cells were acquired using flow cytometry with a FACSanto flow cytometer system and analysed with flowJo, LLC software (BD Bioscience).

### Cell cycle assay

2x10^5^ cells were collected, rinsed with 1xPBS and resuspended in 200 μl of 1xPBS. Cells were fixed adding drop by drop ice-cold absolute ethanol. Fixed cells were washed twice with ice-cold 1xPBS and incubated 1 h at 4 °C in 1xPBS, 25 mg/ml propidium iodide (PI, Sigma, P 4170), 0.5 mg/ml RNase A (Invitrogen, EN0531). After staining, cells were acquired using flow cytometry with a FACSCanto flow cytometer system and analysed with flowJo, LLC software (BD Bioscience).

### The 4 fractions sequential analysis of macromolecules accessibility sequencing (4f-SAMMY-seq), DNA extraction, library preparation and sequencing

The 4f-SAMMY-seq was performed on 5x10^5^ MEFs at 90% confluency using 12 units of DNase I (Invitrogen, AM2222) as described in^42^.

Libraries were then qualitatively and quantitatively checked on and run on the TapeStation System. Libraries with distinct adapter indexes which were normalized to a concentration of 2 nM, equimolarly, pooled, and then loaded onto the Illumina NextSeq 2000 instrument. The sequencing was performed with a minimal target of 15 million reads for 100 bases in single-end mode on the Illumina NextSeq 2000 instrument.

### RNA extraction, library preparation and sequencing

Total RNA was extracted from 1x10^6^ MEFs at 90% of confluence using TRI-Reagent (Sigma, T9424) following the recommended guidelines. The quantification of total RNA was performed with a Qubit 4 fluorometer using the Qubit RNA BR Assay Kit (Invitrogen, Q10210), and RNA integrity was assessed by the Agilent 2100 Bioanalyzer with the Agilent RNA 6000 Nano Kit (Agilent, 5067-1511). For each sample, 10 ng of total RNA was used to construct a strand-specific RNAseq library with SMARTer Stranded Total RNA-Seq Kit - Pico Input (Takara, 634487). The quality and the size of the libraries were analysed with TapeStation System according to the assay guide. The final RNAseq libraries were adjusted to a concentration of 4 nM, equimolarly, pooled, and then loaded onto the Illumina NextSeqTM 550 system. A sequencing depth of 20 million for 75 bases in paired-ends mode was achieved for each sample.

### Chromatin Immunoprecipitation sequencing (ChIP-seq)

MEFs were grown to 90% confluence in DMEM (Gibco, 10566-016) supplemented with 10% (v/v) FBS (Gibco,10270106) and then fixed 10 min at RT with 1% formaldehyde (Sigma-Aldrich, F8775), adding 1:10 of formaldehyde solution (50 mM HEPES-KOH pH 7.5, 100 mM NaCl, 1 mM EDTA, 0.5 mM EGTA, and 11% formaldehyde) on cultured cells. Formaldehyde was then quenched with addition of Glycine (Carlo Erba, 453807) to a final concentration of 125 mM for 5 min at RT, followed by two washes with cold 1xPBS. Cross-linked cells were rapidly collected in a falcon tube by scraping on ice and centrifuged at 2000 g at 4 °C. Pellets of 3x10^6^ cells were stored at -80 °C until sonication. Each 3x10^6^ cell pellet was resuspended in 600 μl of cold Lysis Buffer: 50 mM HEPES-KOH, pH 7.5, 10 mM NaCl, 1 mM EDTA, 10% glycerol, 0.5% NP-40, and 0.25% Triton X-100. After 10 min on a rotator at 4 °C, samples were centrifuged for 5 min at 1350g at 4 °C and resuspended in 130 μl of sonication buffer: 10 mM Tris-HCl pH 8.0, 2 mM EDTA, 0.25% SDS, 1x Protease Inhibitor Cocktail (Roche, 04693116001); 1 mM PMSF (Sigma-Aldrich, 93482). Cells were subjected to lysis on ice for 1 h and homogenised by pipetting every 15 min. Total extracted chromatin was sonicated in a Covaris M220 focused-ultrasonicator using snap cap microTUBEs (Covaris, 520045) (water bath set to 7 °C, peak power 75.0, duty factor 10.0, cycles/burst 250, duration 420 s). Fragmentation of chromatin to an average size of 150-500bp was checked on Agilent 2100 Bioanalyzer using High Sensitivity DNA Kit (Agilent, 5067-4626). To reduce the SDS, samples were dilute to a final concentration 0.1% SDS in equilibration buffer: 10 mM Tris-HCl pH 8.0, 233 mM NaCl, 1.66% Triton X-100, 0.166% Deoxycholic acid sodium salt (DOC), 1 mM EDTA, 1x Protease Inhibitor Cocktail (Roche, 04693116001), and 1 mM PMSF (Sigma-Aldrich, 93482). Samples were then centrifuged at 14000 g for 10 min at 4 °C to pellet insoluble material. Supernatants were quantified using Nanodrop 1000 spectrophotometer and 80 ug of chromatin was used for each immunoprecipitation in a final volume of 300 μl of IP buffer: 10 mM Tris-HCl pH8, 140 mM NaCl, 1.66% Triton X-100, 0.166% DOC, 0.1% SDS, 1 mM EDTA, 1x Protease Inhibitor Cocktail (Roche, 04693116001); 1 mM PMSF (Sigma-Aldrich, 93482). 3% of total chromatin was preserved at 4 °C as input normalization-control for each experimental condition. The remaining chromatin was incubated overnight on a rotator at 4 °C with 6 μl of H3K9me3 (Abcam, ab8898) or H3K27ac (Abcam, ab4729) or H3K4me3 (Sigma-Aldrich, 07-473) antibodies. 20 μl of Protein G beads (Life Technology, 1004D) for each IP were washed twice in 0.1% BSA/IP buffer and incubated on rotator overnight at 4 °C. On the next day, protein G beads were added to each sample and incubated on rotator for 2 hours at 4°C. Beads/IP complexes were then washed 10 min on rotator at 4 °C twice with IP buffer, twice with high-salt IP buffer: 10 mM Tris-HCl pH 8, 500 mM NaCl, 1.66% Triton X-100, 0.166% DOC, 0.1% SDS, 1 mM EDTA, 1x Protease Inhibitor Cocktail (Roche, 04693116001); 1 mM PMSF (Sigma-Aldrich, 93482); twice with RIPA-LiCl buffer: (10 mM Tris-HCl pH 8.0, 1 mM EDTA, 250 mM LiCl, 0.5% DOC, 0.5% NP-40, 1x Protease Inhibitor Cocktail (Roche, 04693116001); 1 mM PMSF (Sigma-Aldrich, 93482); twice with 10 mM Tris-HCl pH 8.0. Crosslinking was reversed by incubating the beads and input at 65 °C overnight with 100 μl of Elution buffer: 10 mM Tris-HCl pH 8, 0,5 mM EDTA, 300 mM, and 0.4% SDS. The next day, all samples were diluted with 100 μl of 1x TE buffer, treated with 2.5 U of RNAse cocktail (Ambion, AM2286) at 37 °C for 120 min, followed by addition of 100 ug of Proteinase K (Invitrogen, AM2548) at 55 °C for 120 min. DNA was then isolated using phenol/chloroform (Sigma-Aldrich, 77617) extraction, followed by a back extraction of phenol/chloroform with additional volume of 1x TE buffer. DNA was precipitated in 2 volumes of cold ethanol, 0.3 M sodium acetate and 20 ug glycogen (Ambion AM9510) for overnight at -20 °C. Pellets were suspended in 31 μl of nuclease-free water and quantified using Qubit 2.0 fluorometer with Qubit dsDNA HS Assay Kits (Invitrogen, Q32854). The libraries were then prepared using the NEBNext Ultra II DNA Library Prep Kit for Illumina (NEB, BE7645L) and NEBNext Multiplex Oligos for Illumina (NEB, BE6440S). Libraries were then qualitatively and quantitatively checked and run on the TapeStation System. Libraries with distinct adapter indexes were normalised to a concentration of 2 nM, equimolarly pooled, and then loaded onto the Illumina NextSeq 2000 instrument. The sequencing was performed with a minimal target of 15 million reads for 100 bases in single-end mode on the Illumina NextSeq 2000 instrument.

### DNA sequence analysis

The results of the sequencing were demultiplexed with bcl2fastq (https://support.illumina.com/sequencing/sequencing_software/bcl2fastq-onversion-software.html, v2.19.0.316). Both 4f-SAMMY-seq and ChIP-seq high-throughput sequencing lanes were merged with the help of GNU parallel software (http://www.gnu.org/s/parallel). All sequencing reads were trimmed using Trimmomatic (v0.39)^74^ with clip file ‘TruSeq3-SE-2. fa’ and the following parameters: 2:30:10 for seed mismatch, palindrome threshold and simple threshold, respectively; 4:15 for sliding window. The minimum threshold of 36 bp length was applied for all reads. Trimmed reads were aligned using BWA (v0.7.17-r1188)^75^ setting -n 2 - k 2 parameters and using as reference genome the mm10 version downloaded from refgenie^76^. The result was saved in BAM file format. PCR duplicates were marked and removed with Picard (v2.23.9) (https://github.com/broadinstitute/picard) ‘MarkDuplicates’ option, collecting the filtered reads in another BAM file. All the reads with mapping quality lower than 1 were filtered out with Samtools (v1.11)^77^ creating another BAM file used for subsequent analyses.

### Genomic reads distribution profiles

For each alignment, coverage analyses were performed using Deeptools (v3.5.0)^78^ with ‘bamCoverage’ function. Original trimmed reads of 100 bp were extended up to 250 bp and normalized with RPKM method with a bin size of 50 bp. The mm10 size was considered of 2652783500 bp, as suggested in the Deeptools manual https://deeptools.readthedocs.io/en/latest/content/feature/effectiveGenomeSize.html) and blacklisted regions, known to be problematic in terms of sequencing reads coverage, were obtained by ENCODE portal (https://www.encodeproject.org/files/ENCFF547MET) and excluded from the analysis. To generate the normalized relative enrichment ratios along the genome for each sample using the 4f-SAMMY fractions, the SPP, R package (v1.16.0)^79^ was used and the library was built under R (v4.1.2). The relative enrichment ratio between 4f- SAMMY-seq fractions was calculated per each biological replicate by comparing the S2S reads against the S3 reads, which was used as baseline 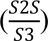. Fraction specific reads were imported from BAM files with ‘read.bam.tags’ function, additionally filtered with ‘remove.local.tag.anomalies’, and the relative differential enrichment was computed using ‘get.smoothed.enrichment.mle’ function setting ‘tag.shift = 0’ and ‘background.density.scaling = TRUE’. The resulting computed signal corresponded to the log2 ratio between the pair of sequencing samples. We defined the solubility profile as the relative enrichment ratio of 4f- SAMMY-seq sequencing reads distribution along the genome for S2S vs S3 fractions. To maximise comparability with 4f-SAMMY-seq data, we computed the normalized relative enrichment ratios of ChIP-seq IP over INPUT genomic profiles using the same methodology described above. To compute correlations between genomic tracks, the smoothed differential signal enrichment was re-binned with Deeptools ‘multiBigwigSummary’ at 50 kb. Genome-wide Spearman correlations between 4f-SAMMY-seq fractions and between 4f-SAMMY-seq solubility profile and ChIP-seq was computed using Deeptools, using the function ‘plotCorrelation’ with the following settings: ‘--corMethod spearman -p heatmap --skipZeros’.

### Enrichment ratio normalization and consensus generation

Each solubility profile was imported with R (v4.3) using the library GenomicRanges (v1.52.0)^80^, resized at genome wide level with 50 kb window. Adjacent blacklisted regions were merged if separated by less than 50 kb with the function ‘reduce’ and setting ‘min.gapwidth= 50000’. It is worth remarking that having the resulting binned genome with a fixed window size of 50 kb, the genomic region adjacent and upstream to the merged blacklist region could be reduced to a different length <50 kb. On the other hand, the genomic region adjacent and downstream to the merged blacklist region were extended to 50 kb, reproducing the original window size. All solubility profiles were normalized by quantile normalization with the preprocessCore library (v1.62.1) (https://doi.org/10.18129/B9.BIOC.PREPROCESSCORE) using the function ‘normalize.quantiles’. The consensus track of each group of samples was generated by computing the mean *x̄* of normalized solubility profiles for each genomic window. The shaded areas represent standard error intervals calculated as 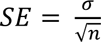 where σ is the standard deviation and n is the number of samples.

### Genomic track representation

The visualisation of genomic tracks was performed with Gviz R library (v 1.44.1)^81^. The ChIP-seq samples were imported using the function ‘import’ of the rtracklayer library and plotted using the function ‘plotTracks’ setting the value ‘windowSize = 1500’ to plot a fine grain profile. Visualization of the 4f-SAMMY-seq consensus tracks were computed using all normalized samples of each group (e.g. WT, LmnaKO, LBDKO) setting the parameter type as ‘a’ and ‘confint’ overlayed using the function ‘OverlayTrack’. Single samples mountain plots were computed by setting the parameter type as ‘polygon”. Extra elements of the panel, chromosome ideogram and relative genome axis, were displayed using the functions ‘IdeogramTrack’ and ‘GenomeAxisTrack’, respectively. The murine cytobands of the chromosome ideogram were downloaded from the UCSC golden path (http://hgdownload.cse.ucsc.edu/goldenpath/mm10/database/cytoBand.txt.gz).

### Differential enrichment analysis of 4f-SAMMY-seq

Differential enrichment analyses were calculated on consensus tracks. Using the WT solubility profile, all the genomic regions with a solubility value outside the arbitrary threshold of ≤ ±0.1 were filtered out. Accordingly, the LmnaKO solubility profile was used as reference for LmnaKO vs. LBDKO comparison. Each genomic bin with a solubility score >+0.1 was considered euchromatin (S2S>S3), while each genomic bin with a solubility score <-0.1 was considered heterochromatin (S2S<S3). For each comparison (WT vs. LmnaKO, WT vs. LBDKO and LmnaKO vs. LBDKO), considering ± 2 standard error intervals for the reference strain, differentially enriched bins were identified as genomic regions where all solubility scores of the compared strain fell outside the interval of consensus of reference (WT or LmnaKO). Candidate bins were then validated with a two tailed z-test with a confidence interval of 0.99, with the function z.testfrom BSDA R library (v1.2.2) (DOI: 10.32614/CRAN.package.BSDA) and adjusted using the Benjamini-Hochberg method for multiple testing correction with the ‘p.adjust’ function and the argument ‘BH’. S2S differentially enriched bins were classified as “S2S UP” or “S2S DOWN” if, with respect to the reference strain, the compared strain increased or decreased its solubility score. S3 differentially enriched bins were classified as “S3 UP” or “S3 DOWN” if, in respect to the reference strain, the compared strain decreased (with more negative numerical values) or increased its solubility score.

### ChIP-seq metaprofile

To compute the average profile over a set of genomic regions of interest (i.e., the meta-profile analysis), average enrichment ratio of single H3K9me3 ChIP-seq replica was plotted over the S3 UP and S3 DOWN regions, together with 1 Mb of upstream and downstream regions. Each genomic region (upstream, S3 UP or DOWN, downstream) was further divided in 600 subregions with R (v4.3) using the library GenomicRanges (v1.52.0)^80^ and the function ‘tile’. The mean signal of IP over the INPUT was measured in each tile for all the regions of the metaprofile with the function ‘mean’ and the ChIP-seq signal was smoothed by ggplot2 (https://ggplot2.tidyverse.org) with the function ‘geom_smooth’ using method = ‘gam’ and formula = ’y ∼ s (x, bs = “cs”).

### RNA sequencing analysis

Sequenced reads were analysed with the pipeline nf-core/rnaseq version^82^. The overall quality of the sequenced reads was assessed using FastQC 2 (v0.11.9) (https://www.bioinformatics.babraham.ac.uk/projects/fastqc/). Reads were trimmed and adapters were clipped by cutadapt (v3.4) (https://doi.org/10.14806/ej.17.1.200). Reads overlapping ribosomal RNA (rRNA) were filtered with SortMeRNA^83^ (v4.3.4), considering all the available databases of rRNA. Reads were then mapped in paired end mode with STAR (v2.7.10a)^84^ on the mouse genome mm10 version. Transcripts were quantified using Salmon^85^ on GENCODE (M25)^77^. Basic gene annotation filtered for transcript with HAVANA characteristics (https://www.sanger.ac.uk/project/manual-annotation/#:~:text=The%20HAVANA%20team%20manually%20annotate,well%20as%20poly%2Dadenylation%20features) and protein coding genes. The full matrix with gene level quantification was created by importing transcript abundances with R (v4.3) library tximport (v1.28.0)^86^ using the function ‘tximport’ and setting the parameter ‘type=“salmon”’. The resulting matrix loaded in R (v4.3) was constructed with genes with a total count greater than 15 in all conditions. The counts were normalised with DESeq2 (v1.40.2), using the median of ratios^87^ and accounting for batch effect using the ‘design = ∼ batch + condition’. RNA-seq samples were grouped in different flow cell batches as follows: Batch 1 contained replicates 3 for LmnaKO and 3 for WT. Batch 2 included replicates 2 for WT, 1-2 for LmnaKO, and 1-2-3 for LBDKO. Finally, Batch 3 consisted solely of replicate 4 of the wild type. The differential expression test was done using the default Wald Test and Benjamini and Hochberg correction for multiple tests, to compute p-values and adjusted p-values, respectively. Differentially expressed genes (DEGs) were identified as those with an adjusted p-value < 0.05 and the absolute value of the | Log2 Fold Change | > 3. Due to high p-values in differential expression results, we also applied the ‘lfcShrink’ function of DESeq2 with type ‘ashr’^88^. The volcano plot showing the DEGs have been done with EnhancedVolcano library (v1.18.0) (DOI: 10.18129/B9.bioc.EnhancedVolcano).

### RNA-seq metaprofile

To compute the metaprofile of the solubility score at differentially expressed genes, the solubility profiles were rebinned at a resolution of 2 kb and normalised using Deeptools (v3.5.0). The merging distance between blacklisted regions was fixed at 5 kb. The 4f-SAMMY-seq solubility score was computed per DEG using the command ‘computeMatrix’ from deeptools scale-regions with setting ‘--beforeRegionStartLength 5000 --regionBodyLength 5000 --afterRegionStartLength 5000 --skipZeros’. The metaprofile was plotted using the command ‘plotProfile’ with the specific settings --plotType heatmap --yMin -0.25 --yMax 0.75’.

### Chromatin compartments analysis

Chromatin compartments were calculated using a revised version of the CALDER algorithm (version 1.0)^89^ as implemented in the original 4f-SAMMY-seq pipeline^42^ . Namely, for each chromosome: i) the four 4f-SAMMY-seq fractions’ (S2S, S2L, S3, and S4) reads distribution profiles were calculated and normalized with RPKM using Deeptools (v3.5.0), rebinned at 50 kb and merged blacklisted regions were filtered out using R (v4.3); ii) for each genomic bin, defined as a vector containing the four RPKM values (one for each fraction), the Euclidean distance (dist, R stats package, method=‘euclidean’) was calculated with all the other bins of the same chromosome. These steps produced an NxN matrix, where N is the number of bins for the considered chromosome. Starting from this matrix, the eigenvector was derived to reconstruct chromatin compartmentalization. Sub-compartment segmentation was limited to the highest level, thus obtaining only 2 compartments corresponding to “A” and “B” compartments. The compartment analysis was performed independently for each chromosome. The consensus of compartment calls across strains was produced by labelling each genomic bin (50 kb) according to the most recurring call across samples: for example, if a bin is labelled as A in at least 2 out of 3 WT samples, that bin will be defined as A in the consensus of controls. The definition of compartment shifts was based on the concordant or discordant compartment classification with respect to WT for each 50 kb genomic bin.

### Gene Ontology (GO)

Gene ontology analysis for both RNA-seq differential expression analysis and compartment switches analysis were performed for Biological Process – BP and Molecular Function – MF using R (v4.3) with gprofiler2 library (v0.2.2) (DOI:10.32614/CRAN.package.gprofiler2) and the GO database id ‘e111_eg58_p18_b51d8f08’, taking as final enriched terms only those with p-value < 0.01, calculated by the function ‘gost’ with correction_method = ‘g_SCS’ (Set Counts and Sizes)^90^.

## Notes

### Competing Interest Statement

The authors have declared no competing interest.

