## Supplemental Material for "The molecular basis of lamin-specific chromatin interactions"

### Extended Data Figures

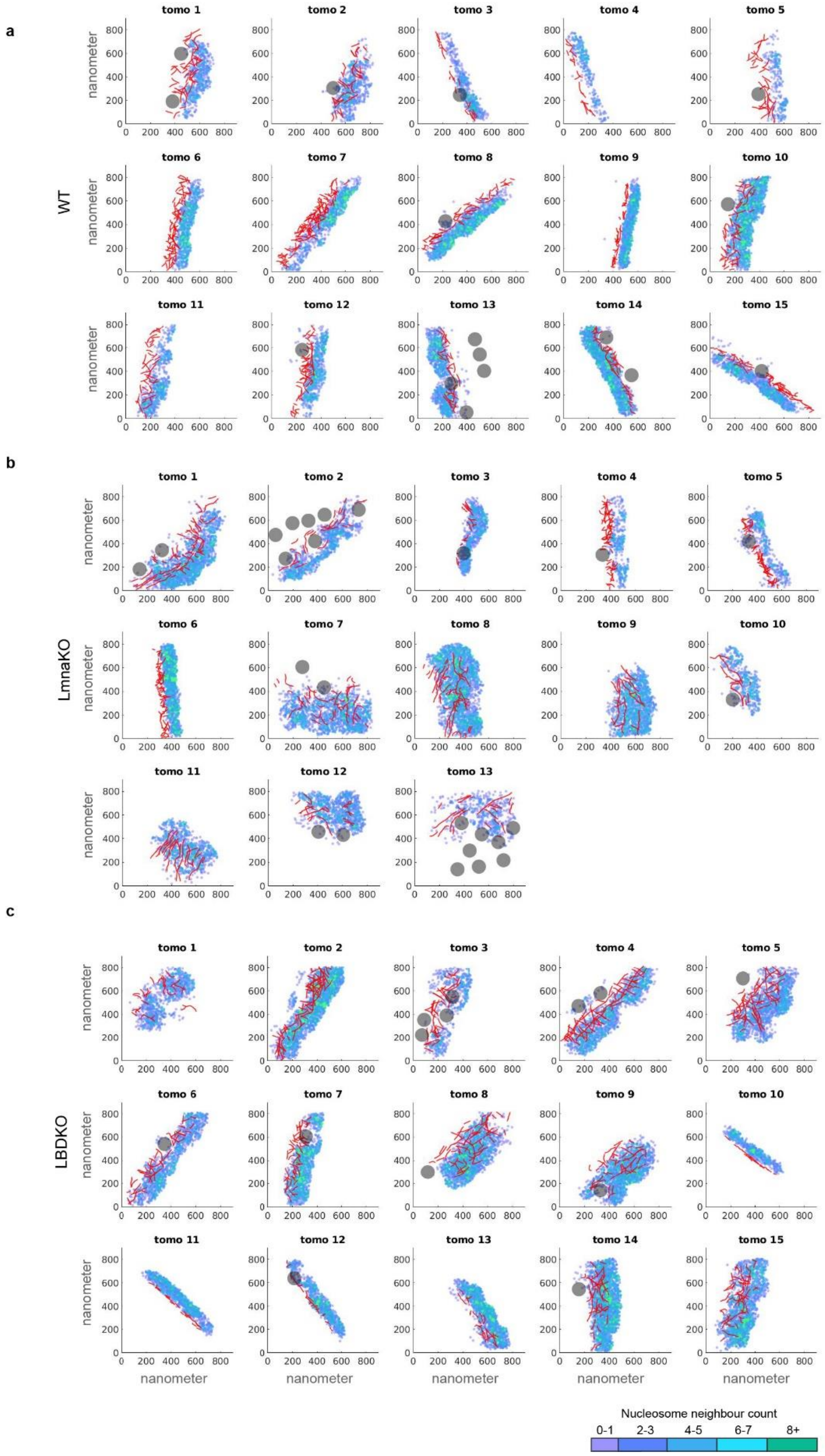

**Extended Data Fig. 1. Gallery of tomograms used for the analysis of WT (a), LmnaKO (b) and LBDKO (c) MEFs.** Each tomogram is depicted by the xy-coordinates of the three analysed structures: lamin coordinates are shown in red, nucleosome coordinates are coloured by increasing concentration, as indicated by the calibration bar, and NPC coordinates are shown in grey.

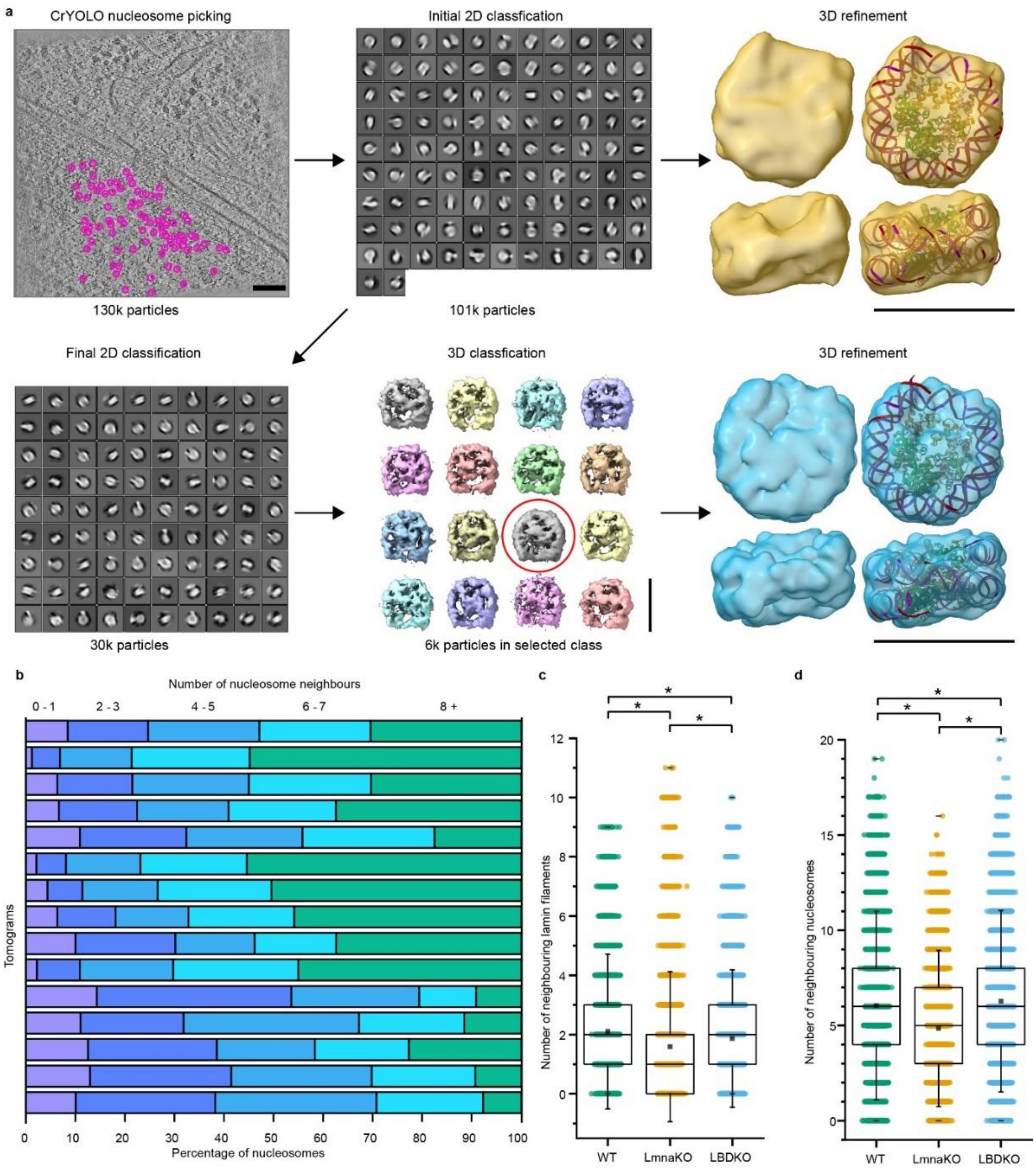

**Extended Data Fig. 2. Overview of nucleosome image processing workflow and measurement of the concentration of nucleosomes and lamins at the NE. a.** In the combined dataset of all three cell lines counting 43 tomograms, an initial set of 130000 nucleosomes were automatically picked by crYOLO[1] , which had been trained and tested based on manual picking of nucleosomes. Upper left panel shows an exploratory result of crYOLO in one tomographic slice of a WT tomogram, scale bar 100 nm. Extracted sub-tomograms were projected and subjected to an initial 2D-classification using RELION[2, 3] to remove false positives. The resulting 101000 nucleosomes were aligned and average in 3D to form a 15 Å structure (yellow), scale bar 10 nm. The coordinates of these aligned particles were the ones used in all further analyses. To improve the structure further 2D- and sequential 3D-classification was performed to obtain a second structure (blue) of 6000 nucleosome at 14 Å resolution, scale bar 10 nm. **b.** The concentration of nucleosomes in each tomogram of WT cells was drawn as a stack bar plot. **c.** For each lamin segment, the number of neighbouring filaments was plotted for each cell line as in **Fig. 2a**. The mean neighbouring filaments count was 2.1, 1.6 and 1.9 for WT, LmnaKO amd LBDKO cells, respectively. **d.** The number of neighbouring nucleosomes for each nucleosome was plotted, resulting in mean values of 6.0, 4.8 and 6.3 neighbouring nucleosomes for WT, LmnaKO amd LBDKO cells, respectively. All boxplots show a box between 25th and 75th percentile, median as a horizontal line, mean as a black square, whiskers represent 1.5 standard deviations. Significance calculated using a one-way ANOVA with a significance level of 0.05. * = $p < 0.05$, n.s. = not significant.

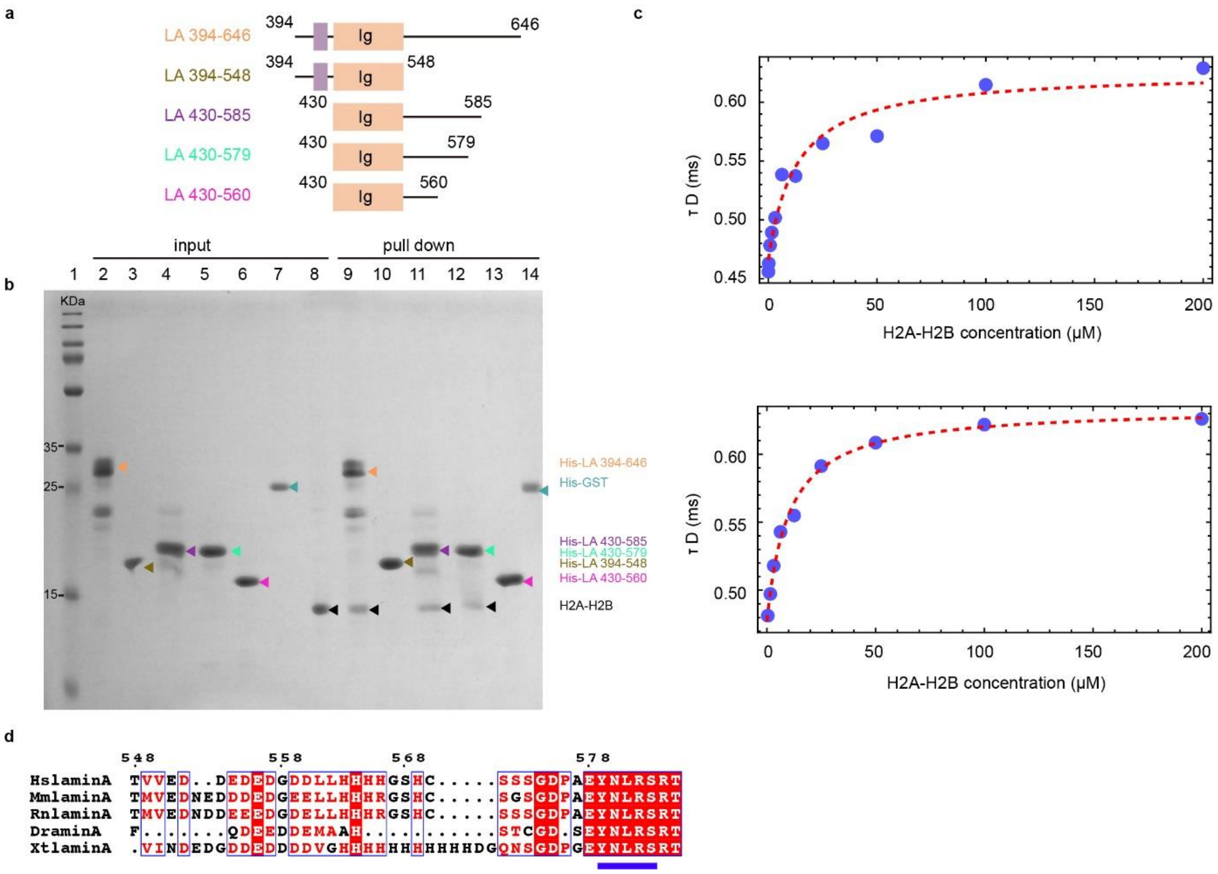

**Extended Data Fig. 3. The lamin A tail domain and its interactions with H2A-H2B. a.** Five truncations of lamin A tail domain were used during in vitro biochemical and structural analyses. **b.** Pull-down assay for identifying the region within lamin A responsible for H2A-H2B binding. 100ug of each candidate truncation, LA 394-646, LA 394-548, LA 430-585, LA 430-579, LA 430-560 and His-GST as a negative control were immobilized on the 200 µl Ni-NTA resin. 5 µl of each sample was analysed by SDS-PAGE (lane 1-6). Purified H2A-H2B heterodimer was shown in lane 7. 100 µg H2A-H2B was applied to each sample and after several rounds of washing and migrated on the SDS-PAGE (Lane 8-13). **c.** Binding isotherms based on the translational diffusion times observed in FCS measurements as a function of lamin concentration yielded a binding affinity between LA 430-585 and H2A-H2B heterodimer of 12 ± 1 µM, confirmed by three biological replicates. **d.** Sequence alignment of lamin A from amino acid 548 to 585 in 5 different species, HslaminA (homosapiens lamin A), MslaminA (musmusculus lamin A), RnlaminA (Rattusnorvegicus lamin A), DrlaminA (Daniorerio lamin A), XtlaminA (Xenopustropicalis lamin A). It shows that the tail domain of lamin A is highly conserved, especially in the region of 578-585 (underlined in blue).

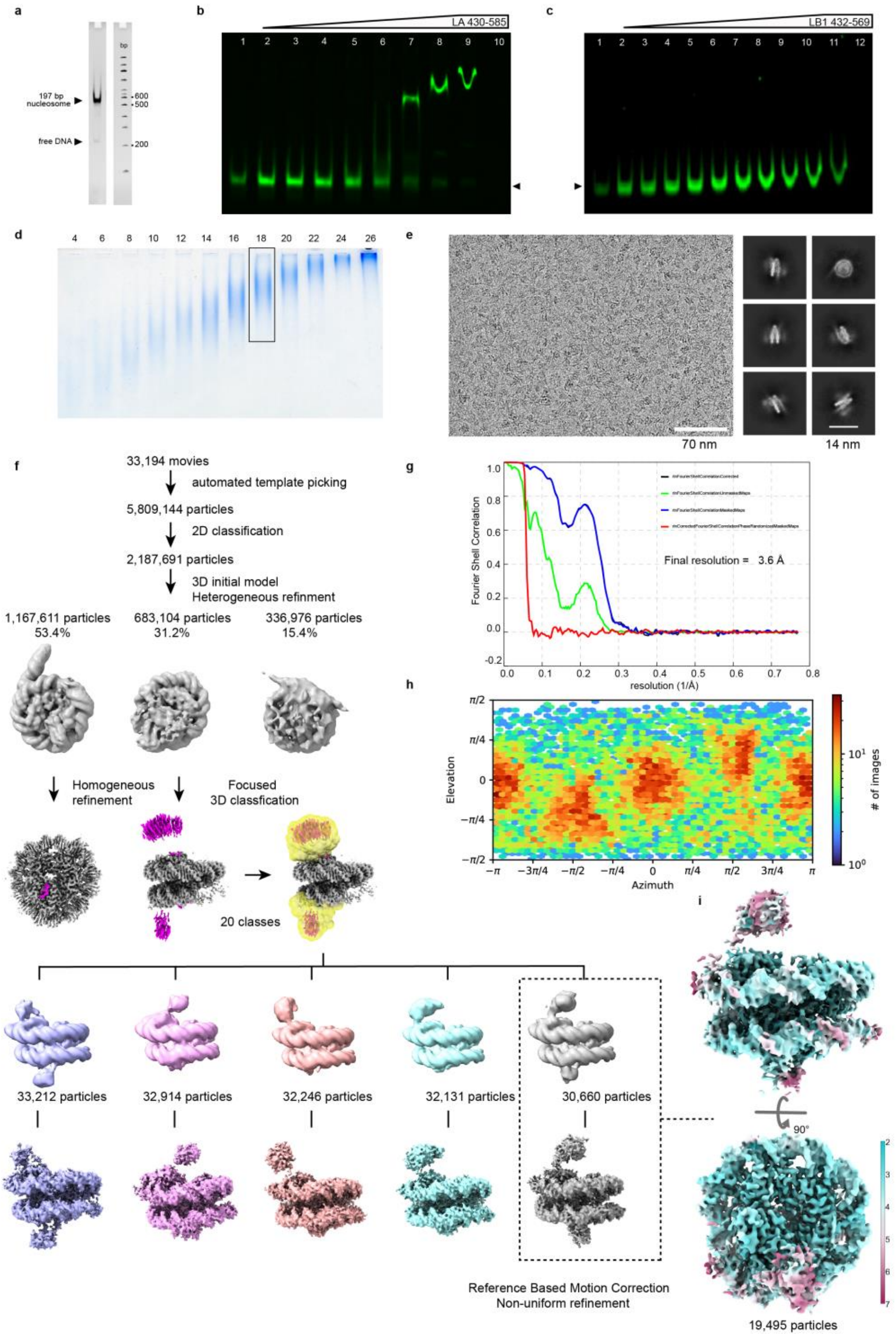

**Extended Data Figure 4. Biochemical and structural analysis of LA 430-585 and nucleosome. a.** In vitro assembled nucleosome was analysed using a 6% native PAGE. **b.** EMSA of LA 430-585 binding to fluorescently labelled nucleosomes. The nucleosome position was marked by black arrowhead. Data was collected from three biological replicates. This experiment was used to quantify the affinity of interactions in **Fig. 3c**. **c.** EMSA of LB1 432-569 binding to fluorescently labelled nucleosomes. There is no shifted band, implying no direct interaction between lamin B1 tail domain and nucleosomes. **d.** LA 430-585 and nucleosome complex after Grafix was analysed by SDS-PAGE. Fraction 18 was collected for cryo-EM sample preparation. **e.** Representative micrograph of the dataset used to determine the structure of LA 430-585 and nucleosome complex. (left) 2D class averages generated from the dataset. (right) **f.** Flow chart for image processing by cryoSPARC. **g.** Gold standard Fourier shell correlation (FSC) curves of final 3D reconstitution (3.6Å). **h.** Angular distribution of particles. **i.** Final reconstruction of the LA 430-585-nucleosome complex coloured by local resolution. Calibration bars are provided.

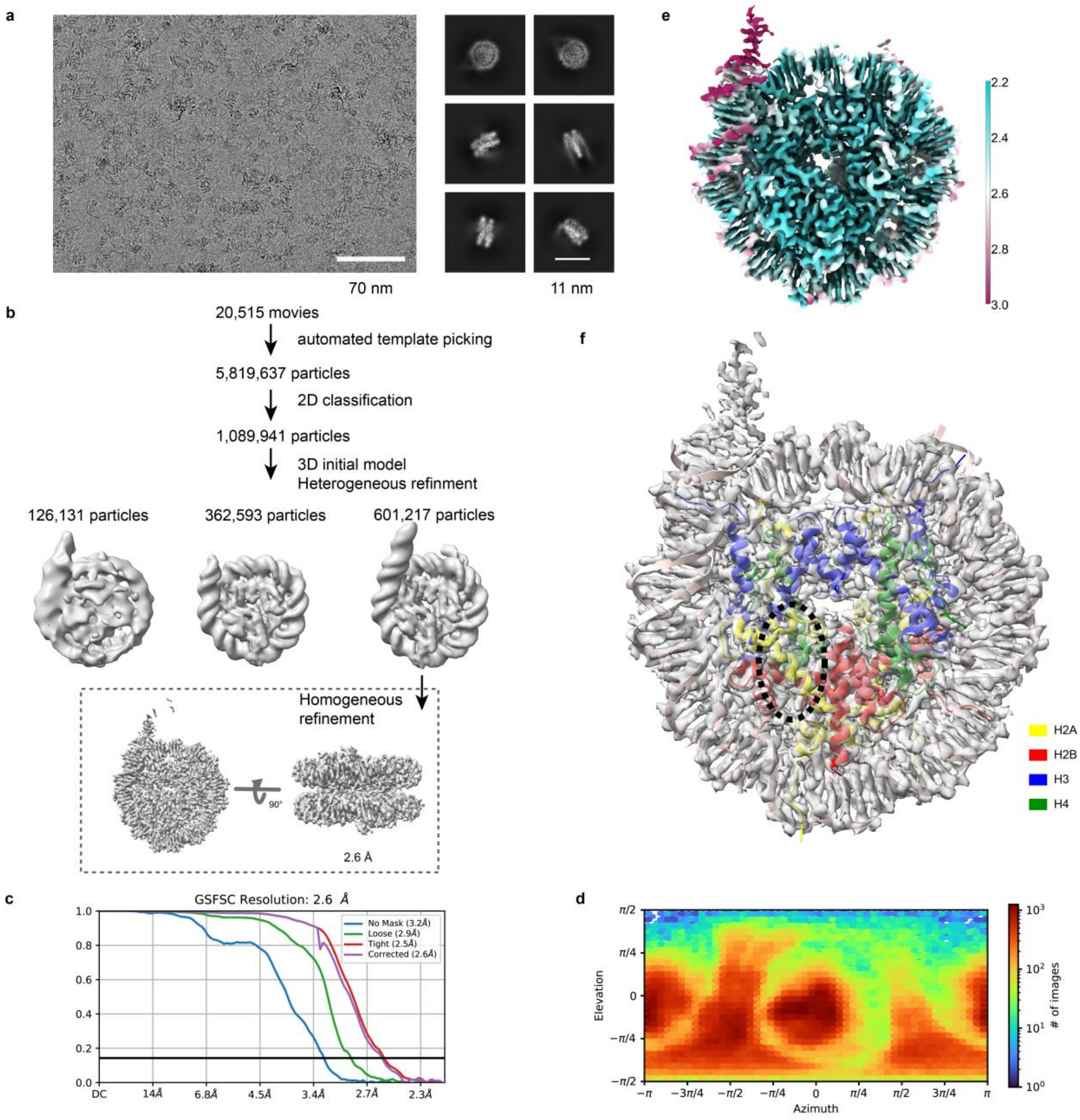

**Extended Data Figure 5. Structural analysis of LA 430-579-nucleosome complex structure. a.** Representative cryo-EM image and 2D class averages of LA 430-579-nucleosome complex. **b.** Flow chart for image processing by cryoSPARC. **c.** FSC curve for final 3D map (2.6Å). Resolution is given for the FSC 0.143 criterion. **d.** Angular distribution of particles. A calibration bar is provided. **e.** Local resolution of final reconstruction. **f.** The final 3D map fitted by nucleosome structure shown there is no more additional density near the acidic patch (black dashed circle).

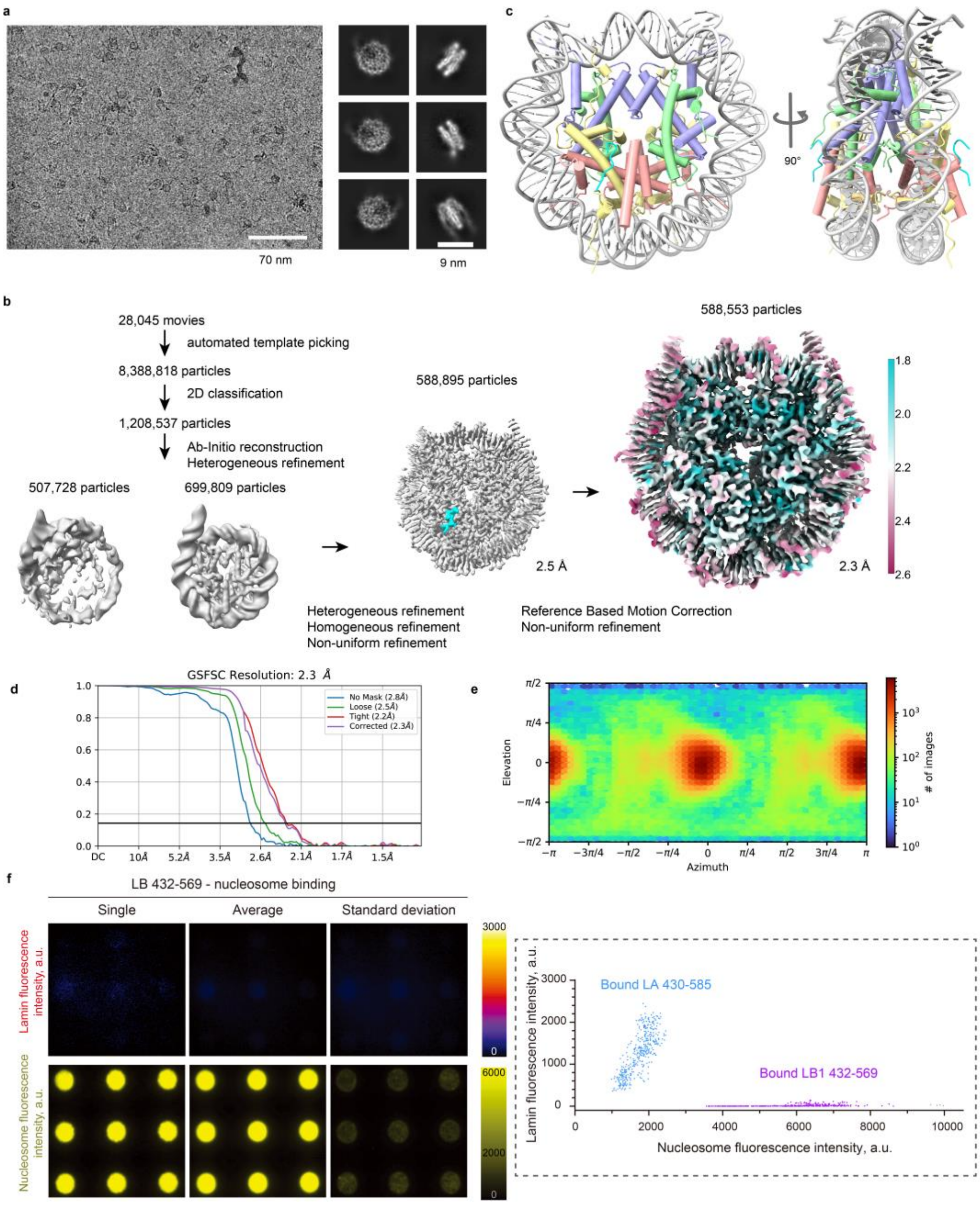

**Extended Data Figure 6. High resolution structural analysis and biochemical binding of peptide (LA 572-588) and nucleosome. a.** Representative micrograph of the peptide-nucleosome dataset was used to determine the complex structure. Typical 2D class averages of the peptide-nucleosome structure. **b.** Flow chart shows the image processing strategy and final 3D map (2.3Å) coloured by local resolution. **c.** Gyre and disc views of the peptide-nucleosome structure shows that peptides are localized on the both sides of H2A-H2B positioned in nucleosome. **d.** FSC curve of final 3D map. **e.** Angular distribution of particles. **b. e.** Calibration bars are provided. **f.** LB1 432-569 does not bind to nucleosomes. Surfaces functionalized with Alexa fluor 488 labelled nucleosomes were incubated with 1 µM of soluble, CF660R fluorescently labelled LB1 432-569 and imaged after 1 hour. The panels show lamin B1 fluorescence intensity (upper row) associated with the spotted fluorescently labelled nucleosomes (bottom row) for a representative single image, for an average image of 18 frames and a total of 162 spots, and its related significantly low standard deviation image. Fluorescence calibration bars are provided. All spots are 3 µm in diameter. The graph on the right shows that nucleosome-associated lamin fluorescence indicated that lamin B1 (Bound LB1 432-569) fluorescence detected on patterned is negligeable compared to that of lamin A (Bound LA 430-585), even for 4 times more immobilised nucleosomes. For lamin A, 399 fluorescence values ranged between 361 and 2446 a.u., and for lamin B1, 370 fluorescence values ranged between 0 and 142 a.u.

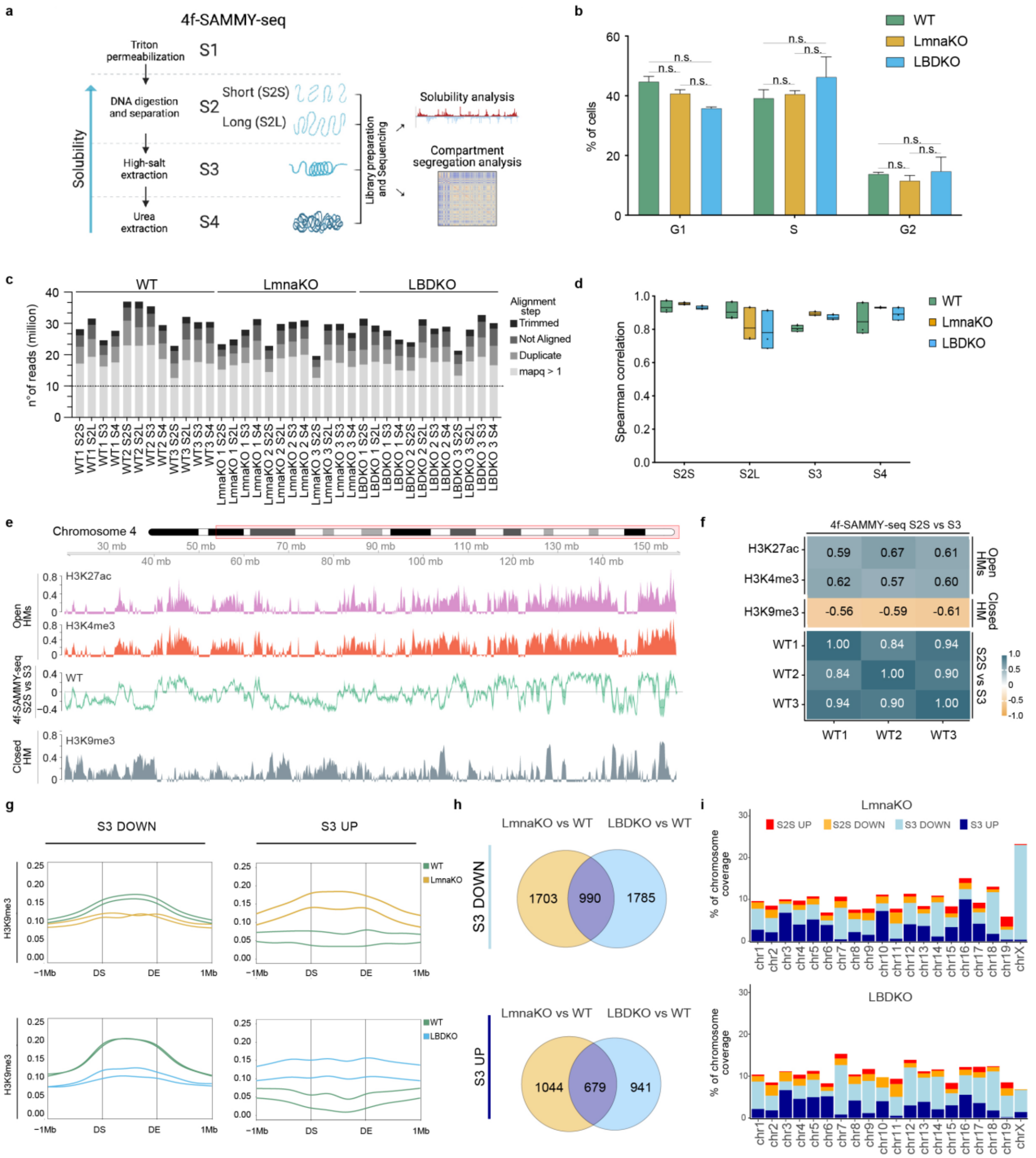

**Extended Data Fig. 7 Euchromatin and heterochromatin distribution captured by 4f -SAMMY-seq. a.** Schematic illustration of the 4f-SAMMY-seq technology. The fractionation process involves the separation of chromatin into distinct fractions: the more soluble DNase-sensitive chromatin (S2), the salt-sensitive chromatin corresponding to compacted heterochromatin (S3), and the salt-resistant chromatin (S4). The S2 fraction is further sub-fractionated based on the size distribution of DNA fragments, resulting in the S2 small and S2 large fractions. Genomic DNA associated with each fraction is extracted, sonicated (only S2L, S3 and S4 fractions), and processed for high-throughput sequencing. Subsequently, the chromatin fractions are mapped to their genomic coordinates, enabling the reconstruction of both euchromatic and heterochromatic regions along with their compartmentalization. **b.** Distribution of cell cycle phases. Fraction of cells in G1, S and G2 phases of the cell cycle in WT (green), LmnaKO (yellow) and LBDKO (light blue). **c.** Stacked bar plots depicting the number of sequencing reads obtained for each sample and chromatin fraction. The bars are divided into trimmed, not aligned, duplicated and mapq >1 (retained for downstream analyses). Dashed, horizontal line represents the minimum of 10 million of mapped reads requested for the analysis. **d.** Boxplot showing the genome-wide Spearman correlation Coefficient (y axis) of each 4f-SAMMY-seq fraction (S2S, S2L,

S3, S4) (x-axis) in WT (green), LmnaKO (yellow) and LBDKO (light blue) replicates. The horizontal lines mark the mean and the boxes extend from the minimum value to the maximum value. **e.** Distribution along a representative region of chromosome 4 (chr4: 20,000,000-END) of 4f-SAMMY-seq consensus track of WT MEF (S2S vs S3). The continuous line represents the mean across the WT replicates and the lighter colour shadows represents the standard deviation. Positive signal enrichments correspond to the more soluble S2S regions, whereas negative signal enrichments represent the less soluble fraction S3. The ChIP-seq tracks for histone mark associated to active chromatin (H3K27ac – purple, H3K4me3 - red), and constitutive heterochromatin (H3K9me3 –blue) are also shown. For ChIP-seq data, the y axis range is set to zero as minimum value. **f.** Genome-wide Spearman correlation for 4f-SAMMY-seq (S2S vs S3 enrichment) between three WT replicates (WT1, WT2, WT3) and histone modifications ChIP-seq (H3K27ac, H3K4me3 and H3K9me3). **g**. Meta-profiles of H3K9me3 enrichment signal over S3 UP and S3 DOWN genomic regions in LmnaKO (top) and LBDKO (bottom). Lines represent two replicates of the H3K9me3 experiment, WT (green), LmnaKO (yellow), and LBDKO (light blue). The x-axis shows the relative position with respect to the start (domain start, DS) and end (domain end, DE) of S3 UP and S3 DOWN genomic regions, along with 1 Mb flanking regions. **h.** Proportional Venn diagram showing the common S3 UP and S3 DOWN genomic regions in LmnaKO and LBDKO. **i.** Distribution of significantly differentially soluble regions across chromosomes in LmnaKO (left) and LBDKO (right) in comparison to WT cells. Colour code: S2S UP (red), S2S DOWN (orange), S3 UP (light blue), S3 DOWN (blue). In **b** the statistical analysis was performed by unpaired two-tailed Student's t-test (ns, p-value > 0.05).

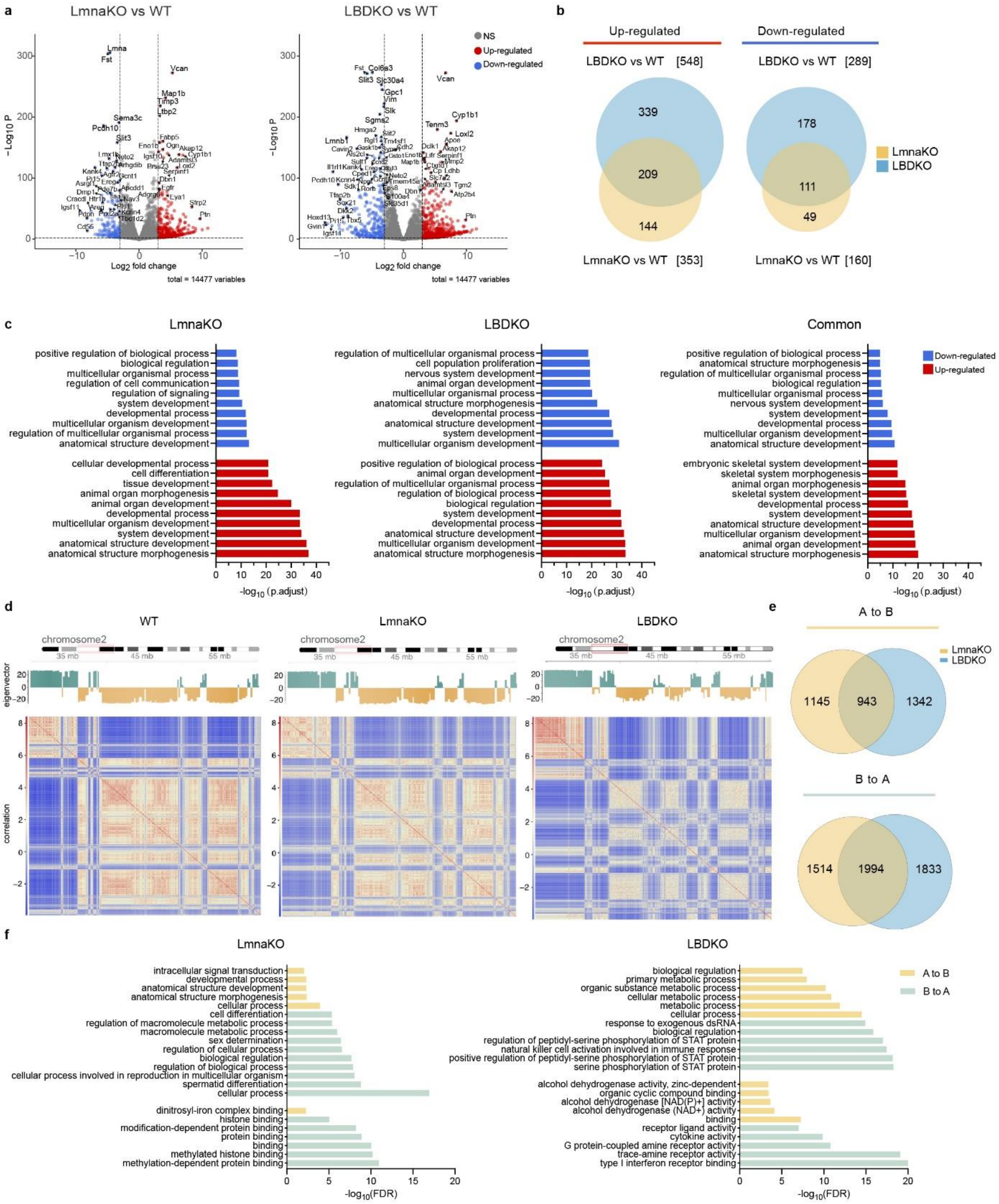

**Extended Data Fig. 8. Lamin specific genome alterations. a.** Volcano plots showing differentially expressed genes (DEGs) between WT and LmnaKO (left), WT and LBDKO (right). The log2FC value > 3, log2FC value < −3 and p-adj < 0.01 (BH) are the cut-off value for significant upregulated (red), and downregulated (blue) Differentially Expressed Genes (DEGs). **b.** Proportional Venn diagram of common up-regulated (left) and down-regulated (right) DEGs in LmnaKO and LBDKO. **c.** Gene Ontology enrichment analysis of upregulated and downregulated DEGs in LmnaKO, LBDKO and common; the bar plots represent the significantly enriched biological processes. **d.** Pair-wise correlation matrices of reads distribution profiles for representative samples WT (left), LmnaKO (middle) and LBDKO cells (right), computed at 50kb genomic bins resolution on chromosome 2. The respective first eigenvector is reported alongside each matrix and coloured to indicate the position of active regions ("A" compartment with positive eigenvalues) and inactive regions ("B" compartment with negative eigenvalues). **e.** Proportional Venn diagram showing common compartments switches ("A to B”) and ("B to A”) in LmnaKO and LBDKO**. f.** Gene Ontology enrichment analysis of genes that change compartment exclusively in LmnaKO (left) or LBDKO (right); bar plots represent significantly enriched biological processes.

**Extended Data Table 1: Cryo-ET data collection and processing of FIB-milled MEFs**

| **Data collection Parameters** | |
|---|---|
| Number of cryo-tomograms used | 43 |
| Microscope | FEI TITAN KRIOS |
| Image detector | GATAN K2 SUMMIT (4k x 4k) |
| Angular range | -60° to 60° (dose symmetric) |
| Tilt increment | 3° |
| Tilts per tomogram | 41 |
| Magnification | 64’000x |
| Voltage (kV) | 300 |
| Electron exposure ($e^-$/$Å^2$) | ~160 |
| Defocus (µm) | -4 |
| Pixel size (Å) | 2.21 |
| **Data Processing** | **Nucleosome structure (EMD-19824)** |
| Number of cryo-tomograms analysed | 43 |
| Pixel size (Å) | 4.42 |
| Initial particles (no.) | 130000 |
| Final particles (no.) | 101780 |
| Symmetry imposed | C1 |
| Defocus (µm) | -4 |
| Map resolution (Å) | 15.8 |
| FSC threshold | 0.143 |

**Extended Data Table 2: Technical details of Cryo-EM structural analysis**

| | Nucleosome-laminA peptide structure (EMD-50114) (PDB-9F0O) | Nucleosome-LA430-585 (EMD-50291) | Nucleosome-LA430-579 |
|---|---|---|---|
| **Data collection and processing** | | | |
| **Magnification** | 130,000x | 130,000x | 130,000x |
| **Voltage (kV)** | 300 | 300 | 300 |
| **Electron exposure ($e^-/Å^2$)** | 70 | 70 | 70 |
| **Defocus range (μm)** | -0.6~-2.4 | -0.6~-2.4 | -0.6~-2.4 |
| **Pixel size (Å)** | 0.65 | 0.65 | 0.65 |
| **Symmetry imposed** | C2 | C1 | C1 |
| **Initial particle images (no.)** | 8,388,318 | 5,809,144 | 5,819,637 |
| **Final particle images (no.)** | 588,553 | 19,495 | 601,217 |
| **Map resolution (Å) 0.143 FSC threshold** | 2.3 | 3.6 | 2.6 |
| **Refinement** | | | |
| **Initial model used (PDB)** | 6ZHX nucleosome | | |
| **Model resolution(Å) 0.5 FSC threshold** | 2.4 | | |
| **Map sharpening *B* factor ($Å^2$)** | -81.7 | | |
| **Model composition** | | | |
| **Non-hydrogen atoms** | 12323 | | |
| **Protein residues** | 790 | | |
| **Ligands** | 0 | | |
| ***B* factor ($Å^2$)** | | | |
| **Protein** | 72 | | |
| **DNA** | 112 | | |
| **R.m.s. deviation** | | | |
| **Bond lengths ($Å^2$)** | 0.013 | | |
| **Bond angles (°)** | 1.84 | | |
| **Validation** | | | |
| **MolProbity score** | 0.5 | | |
| **Clashscore** | 0 | | |

| | |
|---|---|
| **Poor rotamer (%)** | 0.46 |
| **Ramachandran plot** | |
| **Favored (%)** | 98.83 |
| **Allowed (%)** | 1.17 |
| **Disallowed (%)** | 0 |

### Description to Extended Tables

**Extended Data Table 3:** List of differentially expressed genes (DEGs) in LmnaKO and LBDKO MEFs, compared to WT MEFs. In addition, the Gene Ontology (GO) enrichment analysis of molecular function (MF), biological processes (BP), and cellular components (CC) of DEGs that are specific to lamin knockout or commonly deregulated.

**Extended Data Table 4:** Gene Ontology (GO) enrichment analysis of biological processes (BP) and molecular functions (MF) for genes identified in 'A' to 'B' or 'B' to 'A' compartment shifts in LmnaKO and LBDKO compared to WT.
